# A causal spatial role for apical dendrites in the cortex

**DOI:** 10.64898/2026.07.27.741062

**Authors:** Anyi Liu, Kenneth D. Harris, Matteo Carandini, L. Federico Rossi

## Abstract

Cortical pyramidal neurons project a long apical dendrite to layer 1 (L1), to receive synaptic inputs that are thought to provide contextual and possibly suppressive modulation. In visual cortex, these inputs are thought to drive or suppress the responses to large stimuli. Here, we demonstrate a causal role for individual apical dendrites in the awake cortex. We performed two-photon assisted dendritic pruning in individual L5 neurons of the mouse primary visual cortex and discovered that apical dendrites do not suppress the responses to large stimuli; rather, they drive the responses of neurons that prefer such stimuli. To establish the synaptic basis of this effect, we imaged the excitatory synaptic inputs targeting the apical dendrite. We found that these inputs cover an extensive region, which is largest in neurons that prefer large stimuli. These results reveal a clear spatial role for apical dendrites in the cortex.

**Highlights:**

- Optical dendritic pruning in V1 reveals a causal role for L5 apical dendrites
- Apical dendrites drive responses in neurons that prefer large visual stimuli
- Imaging of synaptic inputs to apical dendrites reveals extended apical receptive field
- The apical receptive field is larger in L5 neurons that prefer large visual stimuli

## Introduction

Pyramidal cells, the predominant neurons in the cerebral cortex, send their distinctive apical dendrite upward to form an extensive tuft in layer 1 (L1), where they receive myriad excitatory synapses ^1^. These excitatory synapses arise from neurons in the local cortical region ^2^, in the thalamus ^3^, or in distal cortical areas ^2,4–7^. According to an influential view, these L1 synapses provide top-down signals that are contextual, attentional, or even predictive ^4,8–11^, which the apical dendrite integrates with the putative bottom-up inputs received by basal dendrites. Because the length of the dendrite attenuates the signals ^12^, this integration is thought to rely on active dendritic properties ^9,13–19^.

In addition to excitation, however, the apical tuft also receives inhibition. For instance, the apical tuft receives inhibition from interneurons located in L1 ^20–22^ and in deeper layers ^23– 27^, and the resulting hyperpolarization is visible at the soma ^25,28,29^.

This apical inhibition might play a key role in visual cortex, where it might shape the preferences of neurons for small stimuli. Many neurons in visual cortex are “size tuned”, i.e. prefer small stimuli to larger stimuli ^30–35^. This size tuning is thought to involve inhibition from SST interneurons, which respond to large stimuli ^34,36,37^ and inhibit the apical tuft ^23–27,38^.

It is not known, however, whether in the intact, awake cortex the apical dendrite provides overall drive or suppression to the soma. Activity can be driven through increases in excitation or reductions in inhibition. Similarly, it can be suppressed through increases in inhibition or reductions in excitation. The circuits of the cortex can support all these possibilities. For instance, both NDNF and SST interneurons inhibit the apical tuft, but the latter also inhibit the former ^20^, and are themselves inhibited by other interneurons ^29^.

Distinguishing these possibilities requires targeted measurements and manipulations in the intact, awake cortex. Imaging efforts have revealed the activity of apical dendrites in vivo ^11,39–41^. This activity has typically been found to be coupled with the activity of the soma ^42,43^, with some evidence of compartmentalization ^44^. Efforts to manipulate the apical dendrite in vivo ^10,41,45–48^, in turn, have typically involved entire populations, potentially affecting not only their apical inputs but also the function of the local network. An exception is a groundbreaking study that performed optical apical pruning in individual L2/3 neurons under anesthesia ^49^, but remarkably this study revealed little evidence for a functional role of the apical dendrite, at least in some forms of selectivity.

Here we build on these approaches and establish a causal role for apical dendrites in L5 of the awake visual cortex. We focused on L5 because its pyramidal neurons are the major source of output from cortex ^50^, and we asked whether their apical dendrite is responsible for size tuning, perhaps by suppressing the responses to large stimuli, or possibly for the opposite effect, i.e. driving responses in neurons that prefer large stimuli. We longitudinally recorded from individual L5 neurons in the awake visual cortex before and after optically pruning their apical dendrites ^49^. To establish the synaptic basis of the apical contribution, we then imaged the excitatory synaptic inputs targeting the apical dendrite ^51,52^.

Contrary to our expectations, we discovered that apical dendrites do not suppress the responses to large stimuli in neurons that prefer small stimuli; rather, they drive the responses of neurons that prefer large stimuli. We found that these inputs cover an extensive region, which is largest in neurons that prefer large stimuli. Large-preferring neurons receive apical inputs that cover a larger region of the visual field, are more displaced relative to the receptive field of the neuron, and are more dispersed among each other. These results indicate that apical dendrites preferentially receive inputs from distal regions of cortex, which provide a causal drive to neurons that prefer large stimuli. They thus reveal a causal spatial role for apical dendrites in the cortex.

## Results

To assess the role of apical dendrites in shaping visual responses, we used two approaches. First, we longitudinally recorded from individual L5 neurons in the awake visual cortex before and after optically pruning their apical dendrites ^49^. Then, in a separate cohort of mice, we recorded from the excitatory synaptic inputs ^51,52^ impinging on a neuron’s apical dendrite. We start by describing the first approach.

### Optical dendritic pruning and longitudinal imaging

To investigate the causal contribution of apical dendrites, we established a pipeline to prune these dendrites in single pyramidal neurons in vivo and measure the effects on the activity of the soma. We jointly expressed the red marker tdTomato and the green calcium indicator GCaMP7s in a few L5 pyramidal neurons. To target expression to L5 neurons in the mouse primary visual cortex (V1), we used Rbp4-Cre transgenic mice, which express Cre in L5 excitatory (pyramidal) neurons ^53^. In these mice, we co-injected a diluted Cre-dependent virus expressing Flp and a concentrated virus expressing Flp-dependent tdTomato and GCaMP7s ^54^. We then selected one or more fields of view (FOV, typically 0.6 x 0.6 x 0.6 mm) for longitudinal two-photon imaging. Each FOV typically contained the somas of 12-31 neurons.

In each field of view (FOV) we then followed three longitudinal experimental steps: “Pre” imaging, pruning, and “Post” imaging (Figure 1**a, b**). During the “Pre” session, we imaged the GCaMP7 fluorescence to measure the visual responses of all the neurons in the FOV, and thus established their baseline tuning for stimulus orientation, direction, and size. Mice were awake and head-fixed, but free to run on a wheel (Figure 1**a**). We then acquired a high-resolution structural visualization (Z-stack) to reconstruct the dendritic trees from the tdTomato fluorescence (Figure 1**c**). The following day, during the pruning session, we lightly sedated the mouse (to minimize brain motion) and we randomly selected a subset of visually responsive L5 neurons for two-photon laser pruning of their apical dendrite ^49^. We pruned the apical trunk, aiming right below the main bifurcation, which was typically 150 µm below the cortical surface. The remaining visually responsive neurons in the FOV served as controls. Finally, in the “Post” imaging session, which was typically 1-3 days after pruning, we repeated the same functional and structural imaging steps as in the “Pre” imaging session. This pipeline allows a direct comparison of neuronal functional properties before and after removing apical dendrites, together with a structural confirmation of the apical pruning.

**Figure 1.**
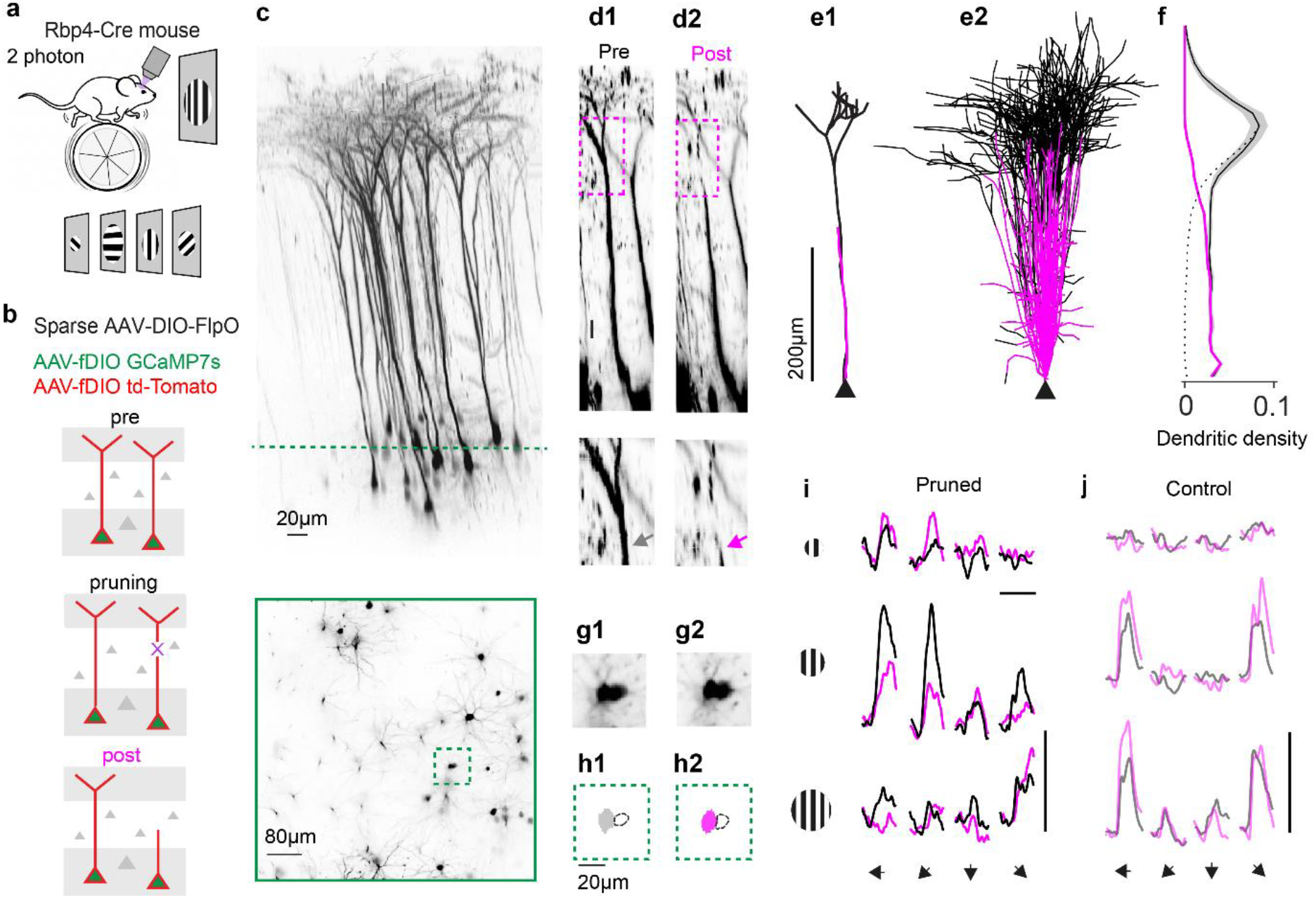
Optical dendritic pruning and longitudinal imaging. (**a**) Schematic of two-photon imaging in V1 and the stimuli used to quantify the visual response properties. (**b**) Schematic of the pruning pipeline, performed in mice transduced with an intersectional viral strategy to sparsely express GCaMP7s and tdTomato in L5 pyramidal neurons. The “Pre” imaging session establishes the neurons’ morphology and baseline visual tuning. In the Pruning session, the apical tufts are pruned in a subset of neurons. The “Post” imaging session repeats the measurements from the Pre session, for comparison. (**c**) Structural imaging of an example FOV. Top: lateral view, showing the somas and the apical dendrites of the neurons in a maximum projection; the dashed line indicates the z position of the section below. Bottom: a horizontal section through the volume, showing the somas and basal dendrites; the dashed square highlights the somas of a neuron selected for pruning and its neighbor. (**d**) Zoomed-in side views showing the apical dendrite of the neuron before (**d1**) and after (**d2**) pruning of the apical tuft. Bottom insets enlarge the site of the pruning. (**e1**) Reconstructions of the neuron in d before (black) and after (magenta) the loss of apical tuft. (**e2**) Same, for all pruned neurons (n=62) recentered by their somatic positions (black triangle), showing overlaid pre (black) and post (magenta) reconstructions. The comparison shows that pruning eliminated all neurons’ apical tufts. (**f**) Marginal distributions of dendritic density along the cortical depth, smoothed and normalized to Pre (black solid line). Dashed line is the difference between the normalized pre and post distributions. (**g**) Mean images of the somas of the pruned neuron in d and a close neighbor, in the Pre (**g1**) and Post (**g2**) session. (**h**) ROI masks extracted from Suite2p for the pruned neuron in the Pre session (**h1**, gray) and in the Post session (**h2**, purple), and for the neighbor (open contours). (**i**) Responses of the example neuron to gratings varying in size (rows) and drift direction (columns), before (black) and after (magenta) apical pruning. Horizontal scale bar, 2 s. Vertical scale bar, 1 normalized response unit, where responses were normalized to the peak response measured within each session. (**j**) Same, for the control neuron in **g** and **h**.

Optical pruning successfully severed the apical tuft while leaving structurally intact the rest of the neuron and the nearby neurons. To measure the effects of pruning, we compared the structural z-stacks acquired in the Pre and Post imaging sessions. Before pruning, a typical neuron showed a vertical apical trunk leading to the arborized apical tuft (Figure 1**d1**). After pruning, the apical trunk showed a clear discontinuity at the targeted site (Figure 1**d2**). To quantify these effects, we semi-automatically traced^55^ the neuron’s morphologies from the soma upwards through the apical trunk. The results demonstrated successful severing of the apical tuft, both in the example neuron (Figure 1**e1**) and in the full cohort of 62 neurons (in 10 FOVs from 5 mice, Figure 1**e2**). Cutting the apical tufts eliminated the dendritic density above the pruned section (Figure 1**f**).

The pruned neurons remained healthy and visually responsive. The soma and basal dendrites of the pruned neurons were unaffected by pruning, as were those of nearby, or even adjacent, neurons (e.g. Figure 1 **d,g,h**). Moreover, the pruned neurons showed healthy responses to visual stimulation both before pruning and after (e.g. Figure 1**i**). These observations indicate that pruning caused no detrimental side effects. We could then concentrate on the visual responses of the neurons and establish the effects of apical pruning, using the non-pruned neurons as a local control.

To study the effects of apical pruning on spatial integration, we divided the neurons into two groups based on their pre-pruning responses to gratings of different sizes. As expected ^30–35^, many neurons responded more strongly to small stimuli than to larger stimuli. A typical neuron among these gave the strongest responses to stimuli with a 20 deg diameter (Figure 2**a**). This property was common: 17 of the 32 neurons that were subsequently pruned preferred small stimuli (5-20 deg diameter, “small-preferring”). The remaining neurons (“large-preferring”) preferred larger stimuli (40-80 deg diameter).

**Figure 2.**
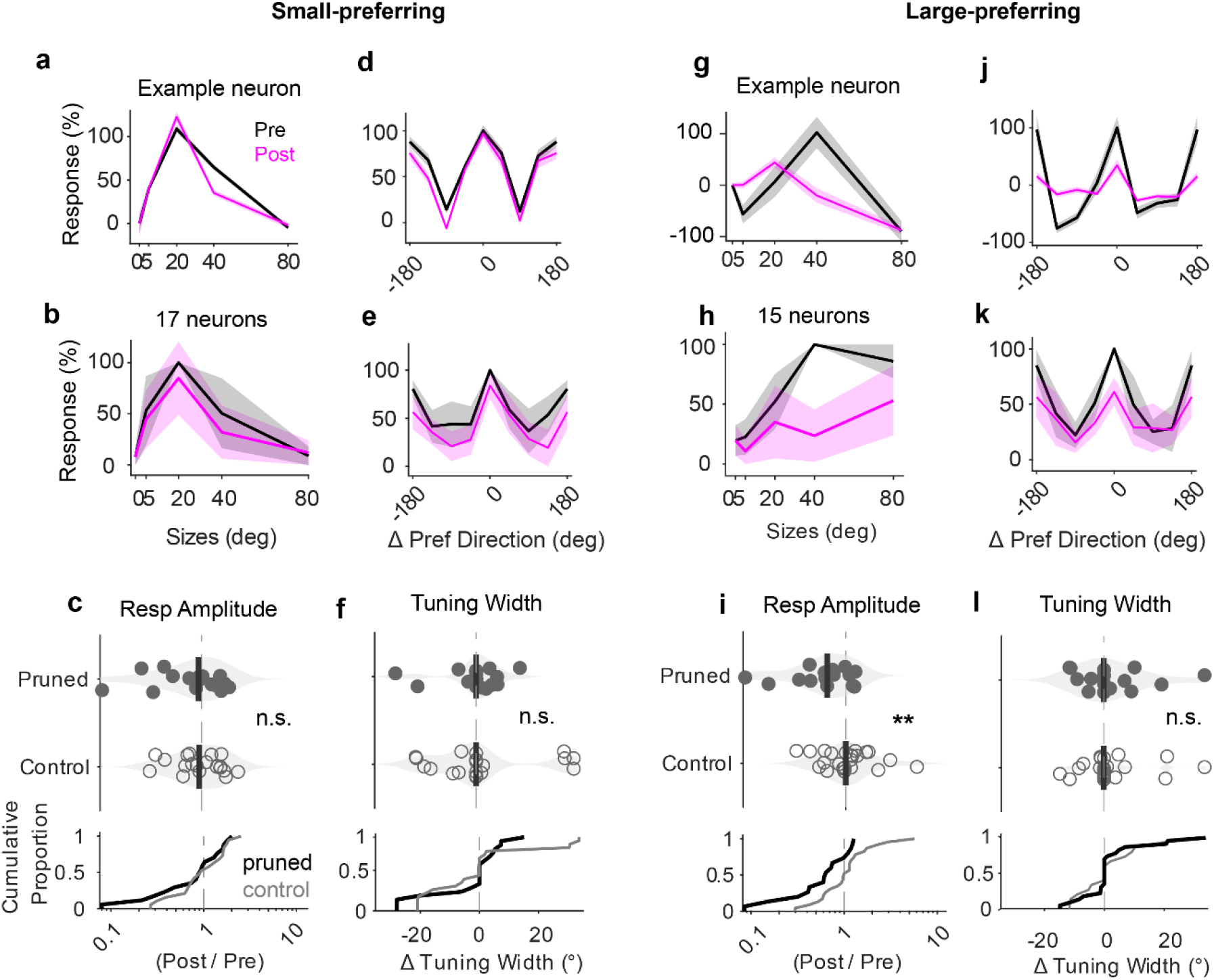
Apical dendrites drive visual responses in large-preferring neurons. (**a**) Tuning for stimulus size of an example small-preferring neuron (neuron 31), showing the mean responses before (black) and after (magenta) apical pruning. At each size, the responses are averaged over stimulus orientations and directions, and normalized so that 0 is the response to a blank screen (measured separately in Pre and Post sessions) and 100% is the response to the best stimulus (measured in the Pre session). Shaded regions show ± s.e. (standard error) across repeats. (**b**) Same, for the population of pruned small-preferring neurons (those giving strongest responses to 5 or 20 deg stimuli, n=17). Curves show median values, and shaded areas show ± 1 m.a.d. (median absolute deviation) across neurons. (**c**) Effect of pruning on responses at the preferred size, for pruned small-preferring neurons (n=17, filled dots and lack line) and control small-preferring neurons (n=19, hollow dots and grey line). The x axis shows the ratio of responses in Post session relative to the Pre session. (Median effect: control vs pruned small-preferring neurons: 0.943 vs 0.927, p = 0.181, (n.s.) Permutation test). (**d**) Same as **a**, showing the example neuron’s tuning for stimulus orientation and direction. At each orientation and direction, the responses are averaged over all sizes. (**e**) Same as **b**, showing the population’s tuning for stimulus orientation and direction. (**f**) Same as **c**, showing the effect of pruning on orientation tuning width. (Median effect: control vs pruned small-preferring: -1.98e-08 vs 0.116, p = 0.334, (n.s.) Permutation test). (**g**-**l**) Same as **a-f**, for neurons preferring large stimuli (40 or 80 deg), showing an example neuron (**g**,**j**) and the population of pruned neurons (n = 15, **h, k**). (**i**: effect on preferred size response, median effect: control vs pruned large-preferring neurons: 1.00 vs 0.636, p = 0.007, (**) Permutation test; **l**: effect on preferred orientation tuning width, median effect: control vs pruned large-preferring neurons: -3.45e-09 vs -1.59e-08, p = 0.439, (n.s.) Permutation test).

### Apical dendrites do not influence visual responses in small-preferring neurons

We first focused on neurons selective for small stimuli, and found their responsiveness and size selectivity to be immune to apical pruning. The example small-preferring neuron was not noticeably affected by the loss of the apical dendrite (Figure 2**a**). Similar effects were seen in the population of small-preferring neurons (Figure 2b). After pruning, their responses showed no more variation than those of unpruned neighboring control neurons (Supplementary Figure 1; p = 0.181, permutation test, Figure 2**c**).

This robustness to apical pruning extended to these neurons’ tuning for stimulus orientation and direction. The tuning for orientation and direction remained similar before and after apical pruning, both in the example neuron (Figure 2**d**) and in the population of small-preferring neurons (N = 15, Figure 2**e**). The orientation tuning width showed no more variation in pruned neurons than seen in control neurons (p = 0.727, permutation test, Figure 2**f**). Similar results were seen for direction selectivity (p = 0.725, Supplementary Figure 2).

These results indicate that the apical dendrite does not contribute to the visual responses of small-preferring neurons. In these neurons, the apical dendrite does not seem to amplify or suppress the responses, nor does it participate in these neurons’ preference for small stimuli. Moreover, consistent with previous results obtained in L2/3 under anesthesia ^49^, the apical dendrite is not required to establish the neurons’ selectivity for stimulus orientation and direction.

### Apical dendrites drive visual responses in large-preferring neurons

In contrast, apical pruning reduced visual responses in large-preferring neurons. A typical neuron among these gave the strongest pre-pruning responses to a 40 degree stimulus, but these responses were drastically reduced following pruning (Figure 2**g**). A similar effect was seen in the population of large-preferring neurons (n=15), which showed a clear reduction in firing rate after pruning (Figure 2**h**). This reduction was significantly larger than observed in the control neurons (p = 0.007, permutation test, Figure 2**i**).

This reduction in responsiveness was not accompanied by changes in orientation and direction tuning. In the example neuron, apical pruning reduced the responses but did not change the overall tuning for orientation and direction (Figure 2**j**). Similar results were seen in the population (Figure 2**k**). After pruning, the orientation tuning width showed no more variation in pruned neurons than in control neurons (p = 0.979, Figure 2**l**). Similar results were seen for direction selectivity (p = 0.210, Supplementary Figure 2).

These results indicate that the apical dendrite is critical for the responses of large-preferring neurons, providing a prominent excitatory drive to the neuron’s responses, particularly to large stimuli.

### Apical inputs extend beyond a neuron’s receptive field

A possible explanation for the results of pruning is that the apical dendrite of large-preferring neurons integrates excitatory inputs from a large region of visual space, i.e. from presynaptic neurons with dispersed receptive fields covering a wider region than small-preferring neurons. Together with receiving inputs from a wider region of cortex, the apical dendrite might also receive more inputs from higher visual areas ^2,5,7^, which tend to have larger receptive fields ^56^ and a preference for large stimuli ^57^. By integrating them, the apical dendrite could thus provide a spatially distributed drive to the soma.

To test this hypothesis, we established a dualcolor imaging approach to record the visual responses of a neuron’s soma and of its apical synaptic inputs. We co-expressed the red calcium indicator jRGECO1a and the green glutamate sensor iGluSnFR4s in a sparse population of L5 neurons (Figure 3**a**). For this co-expression, we used two strategies: (1) in Rbp4-cre mice we injected with a diluted Cre-dependent flp virus and concentrated flp-dependent iGluSnFR, and jRGECO1a virus; (2) in wild-type mice we injected a diluted Cre virus with concentrated Cre-dependent iGluSnFR and jRGECO1a, focusing the imaging on L5 neurons identified by morphology and somatic depth. We could then record the activity of presynaptic neurons synapsing on apical dendrites by imaging iGluSnFR in the apical tuft, and of the postsynaptic neurons by imaging the soma (in a separate acquisition) with jrGECO1a.

**Figure 3.**
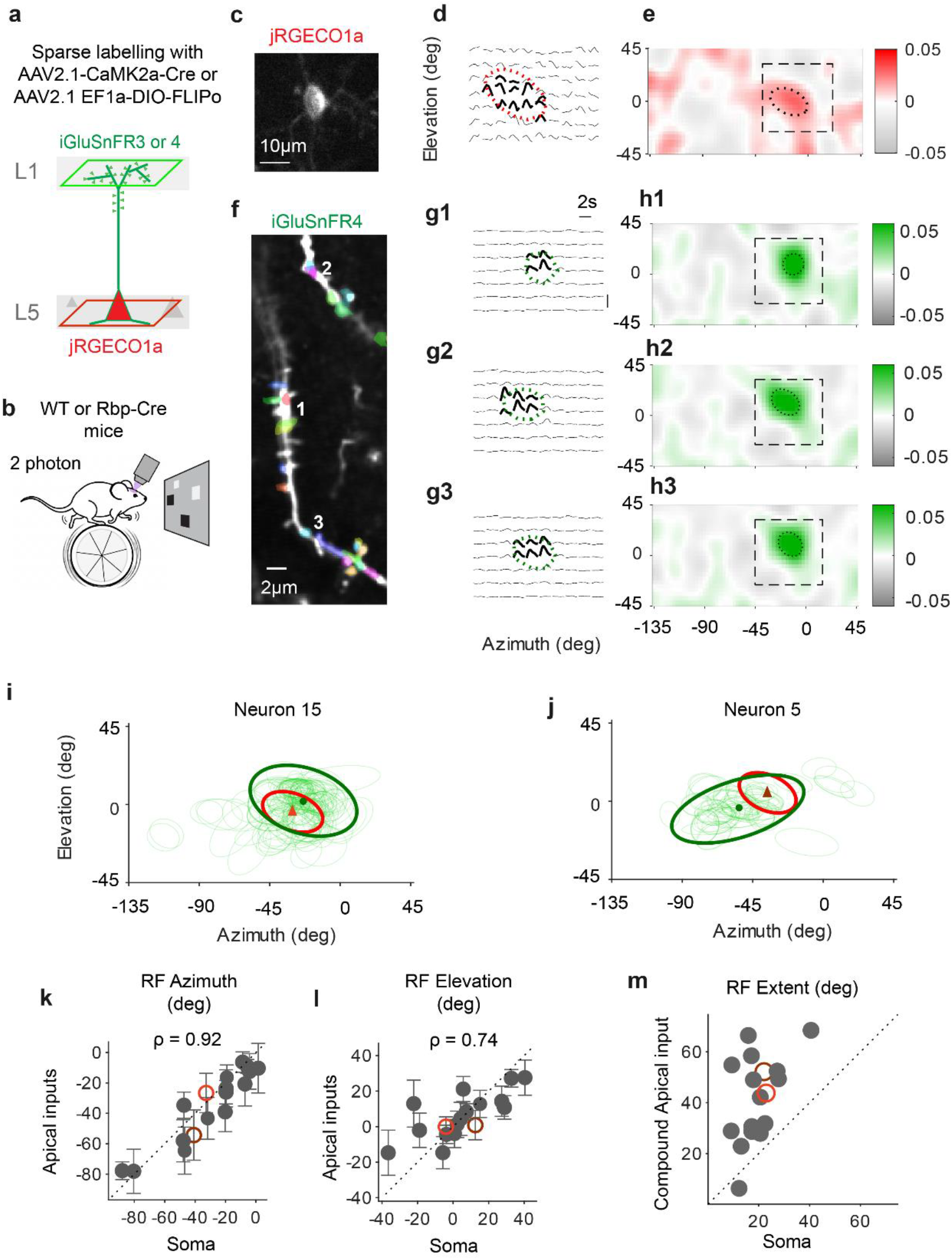
Apical inputs extend beyond a neuron’s receptive field. (**a**) Strategy for dual-color imaging. A few V1 L5 pyramidal neurons are labelled with iGluSnFR (version 3 or 4) for apical dendritic imaging (in L1) and with jRGECO1a for somatic imaging (in L5). (**b**) Receptive fields (RF) are mapped with a sparse noise stimulus. (**c**) Mean image of the soma of an example neuron, from the jRGECO1a signal. (**d**) Normalized responses of the soma to the appearance of a square, as a function square position and time. A 2D Gaussian is fit to the spatial profile and the region containing 40% of its strength is outlined (red ellipse, thick traces). (**e**) Smoothed spatial profile of the responses, interpolated at 1 deg resolution. Square and ellipse indicate the regions in d. (**f**) Mean image of example apical branches of the same neuron imaged in L1 from the iGluSnFR signal. Synaptic ROIs are in arbitrary colors. Numbers indicate the 3 synaptic sites analyzed in **g** and **h**. (**g**) Responses of those synaptic sites as a function of position and time. Format as in d. (**h**) Smoothed spatial profile of the responses. Format as in e. (**i**) RFs of all imaged apical synaptic inputs (thin green ellipses) and of the soma (red ellipse) for the example neuron in **c**-**h**. The thick green ellipse denotes the 90% confidence region of the entire synaptic RF population, measured using a robust covariance estimator. It estimates the RF of the compound apical input. Its center (green dot) is near the center of the somatic RF (red triangle). (**j**) Same, for another example neuron. (**k-l**) Alignment between the RF centers of the soma and of the compound apical input in visual azimuth (**k**, ρ = 0.92, Spearman correlation) and elevation (**l**, ρ = 0.74). Each point represents one neuron; the orange and the dark red circles denote the example neurons in **i** and **j**. Error bars indicate standard deviation in synaptic RF centroid estimation. (**m**) Extent of the compound apical input RF vs the somatic RF. Each point represents one neuron.

These methods enabled us to map the receptive fields of L5 neurons and of the presynaptic inputs that land on their apical dendrites. To map receptive fields (RFs), we first presented sparse noise stimuli with randomly occurring black and white 6 deg wide squares (Figure 3**b**).

First, we imaged the soma using red signals from jrGECO1a (Figure 3**c**), and mapped the neuron’s RF using regularised regression, and summarized its spatial footprint by fitting an ellipse (Figure 3**d, e**). We then moved the imaging plane up to individual apical tuft branches in Layer 1 and increased the sampling rate from 5-6 Hz to 30 Hz, to image the glutamate released at individual dendritic synapses using green signals from iGluSnFR (Figure 3**f**, Supplementary Figure 3). Their visual responses stood out (Figure 3**g**) and were restricted to well-localized RFs, which we also summarized by fitting ellipses (Figure 3**g**,**h**).

A neuron’s RF was generally located close to the centroid of the inputs synapsing on its apical dendrite. By plotting the ellipses corresponding to the RFs of the neurons synapsing on the apical dendrite, we could observe their scatter and summarize it by fitting another, larger ellipse (e.g. Figure 3**i**). This larger ellipse describes a compound apical RF, which typically encompasses the RF of the neuron measured at the soma (Figure 3**i,j**). Indeed, the centers of the somatic RF and of the compound apical RF were highly correlated across neurons, both in azimuth (p < 0.001, N = 17, Spearman correlation, Figure 3**k**) and in elevation (p = 0.001, Figure 3**l**).

In practically all neurons, however, the region covered by the apical input was markedly wider than the neuron’s RF (p < 0.001, Wilcoxon Signed-Rank Test, n = 17 neurons, Figure 3**m**, Supplementary Figure 4). To summarize the extent spanned by somatic and compound RFs we averaged the major and minor axes of their ellipse fit. While the somatic RF spanned 20±8 deg (range 9 to 41 deg), the extent of the compound apical RF was wider (42±17), and varied substantially across neurons (range 6 to 68 deg, Figure 3**m**).

Intriguingly, the extent of the compound apical RF was not related to the extent of the somatic RF (p = 0.171, Spearman’s Rank test, ρ = 0.35, Figure 3**m**, Supplementary Figure 5). Nevertheless, classical RF measured with the sparse noise do not capture the full integration area of a neuron, and may not to correspond to its size preference as determined by gratings of varying sizes ^58,59^. We therefore asked whether the extent of apical compound RFs covaried with the size tuning of somatic responses.

### Apical inputs cover a wider region in large-preferring neurons

In addition to measuring each soma’s RF with sparse noise, we also measured its size tuning with drifting gratings (e.g. Figure 4**a**). As expected, the somatic responses were selective for smaller gratings in some neurons (e.g. Figure 4**a**) and for larger gratings in others (e.g. Figure 4**b**). As before, we split the neurons into two cohorts (Supplementary Figure 4): smallpreferring neurons (n = 8 somas preferring 5 deg or 20 deg stimuli, Figure 4**c**) and large-preferring neurons (n = 9 somas preferring larger stimuli, Figure 4**d**). These size preferences were not related to the extent of somatic RFs (Supplementary Figure 5).

**Figure 4.**
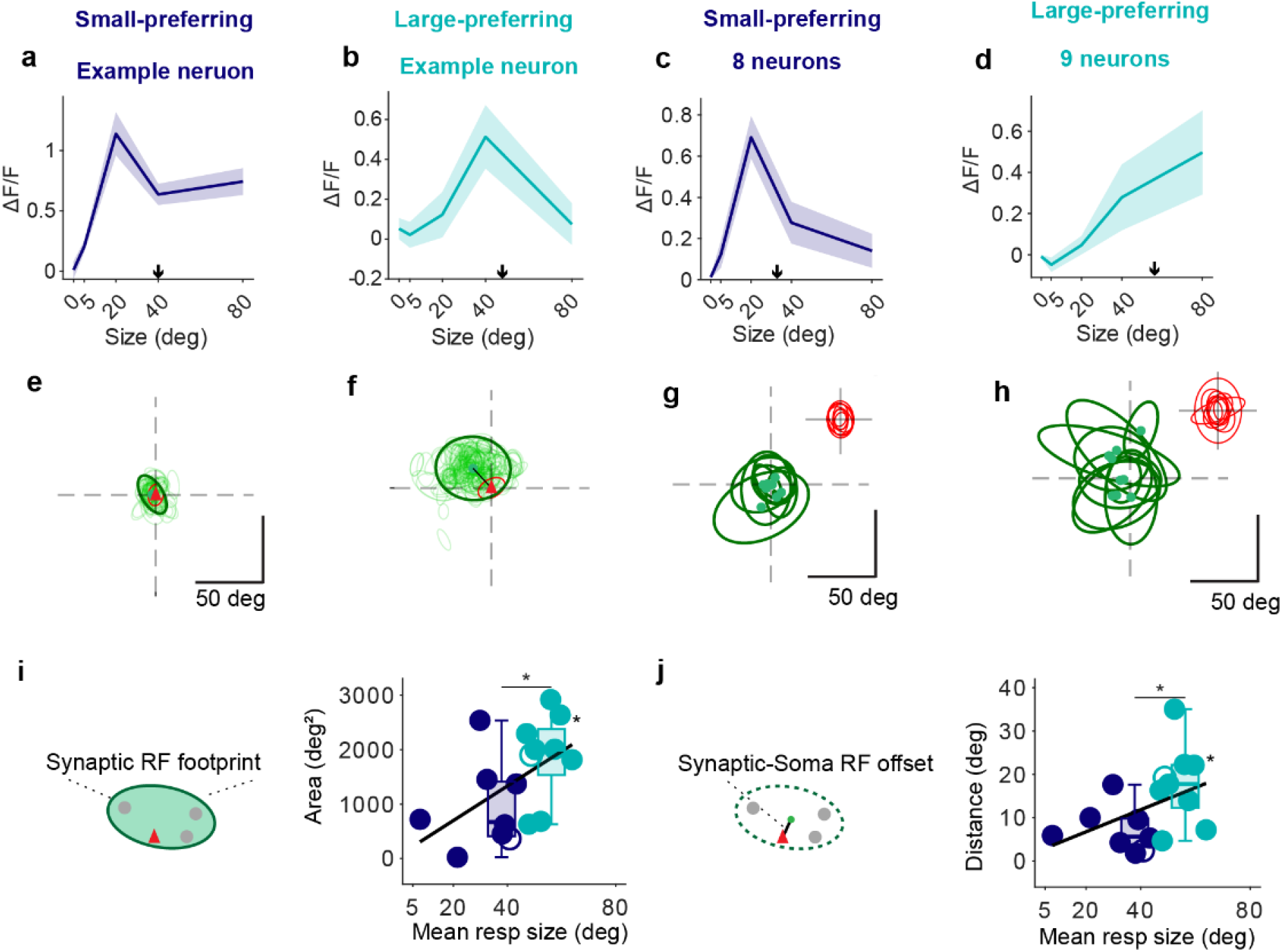
Apical inputs cover a wider region in large-preferring neurons. (**a**) Somatic size tuning curve of an example small-preferring neuron. Shaded area indicates the standard error across stimulus repeats. The center of mass of the responses (arrow) constitutes the mean responsive size. (**b**) Same, for an example large-preferring neuron. (**c**-**d**) Population median size tuning curve. Shaded area indicates standard error across all neurons. (**e**) Apical RFs (thin green ellipses) for the example small-preferring neuron in **a**, relative to the center (red triangle) of the somatic RF (red ellipse). The compound apical RF (thick green ellipse) is the 90% confidence region for the apical RFs. (**f**) Same, for the example large-preferring neuron in **b**. (**g**) Compound apical RFs (green ellipses) of all the small-preferring neurons (n=8), relative to the center of the somatic RF (origin). The somatic RFs are plotted in the inset (top right). Green dots indicate the center of each compound apical RF; black lines show the offset vectors from the soma. (**h**) Same, for large-preferring neurons (n=9). (**i**) Area of the compound apical RF (area of the ellipse) vs mean responsive size for small-preferring (purple) and large-preferring (teal) neurons. Each dot represents a neuron (hollow dots: example neuron from **a, b**), overlaid with group boxplots. Lines indicate linear regression across all neurons. Asterisks indicate slope significance and between-group differences (Wilcoxon rank-sum test; *p < 0.05, **p < 0.01, n.s., not significant). (**j**) Same, for the offset between the compound apical RF and the center of the somatic RF.

Consistent with our hypothesis, the compound apical RF tended to be more compact in small-preferring neurons than in large-preferring neurons. For instance, in the example small-preferring neuron, the RFs of the individual apical inputs clustered tightly around the RF of the soma, thus defining a compound apical RF that was compact (23 deg in extent, Figure 4**e**). By contrast, in the example large-preferring neuron the RFs of apical inputs appeared more distant from the RF of the soma, and more widely dispersed (50 deg in extent, Figure 4**f**). This distinction was confirmed at the population level: the RF of the compound apical input tended to be more compact in small-preferring neurons (extent: 32±16 deg, mean ± s.e. Figure 4**g**) than in large-preferring neurons (extent: 51±14 deg, Figure 4**h**).

Indeed, the compound apical RF was significantly larger in large-preferring neurons than in small-preferring neurons (total area: 1987±309 deg^2^ vs. 670±475 deg^2^, median ± m.a.d., p = 0.027, Wilcoxon Signed-Rank Test, Figure 4**i**). In fact, the area of the compound apical RF correlated with the size tuning of each neuron in a graded manner. For each neuron, we obtained this graded measure by computing the center of mass of the responses (“mean responsive size”, arrows in Figure 4**a-d**). This measure correlated positively with the area of the compound apical receptive field (ρ = 0.50, p = 0.042, Figure 4**i**). This effect did not depend on the variability in number of synaptic inputs sampled per neuron across groups (2-ways ANOVA, p = 0.030 for large vs small neurons, p = 0.60 for synapse count). Consistently, compound RF size did not correlate with synapse count across individual neurons (Spearman’s rank correlation: ρ = -0.075, p = 0.78; Supplementary Figure 6).

In large-preferring neurons, moreover, the compound apical RF tended to be more offset from the center of the somatic RF. For instance, in the example large-preferring neuron the center of the compound apical RF appears markedly displaced from the center of the somatic RF (Figure 4f), much more so than in the example small-preferring neuron (Figure 4e). This effect was confirmed across the population: the offset between the compound apical RF and the somatic RF was larger in large-preferring neurons (18±4 deg vs. 6±3 deg, median ± m.a.d., p = 0.01, Figure 4**j**) and correlated positively with the mean responsive size of the neurons (ρ= 0.50, p = 0.042 Figure 4**j**). This offset was independent from the variability in number of synaptic inputs sampled per neuron across groups (2-ways ANOVA, p = 0.016 for large vs small neurons, p = 0.53 for synapse count), and did not arise from the differences in the compound size (Monte Carlo zero-offset null test, p = 0.0001). As a consequence of these two phenomena, the apical inputs to large-preferring neurons were also more dispersed relative to each other and to the neuron’s somatic RF (Supplementary Figure 6).

The responses to stimuli of different size, therefore, exposed a simple logic in the arrangement of synaptic inputs to the apical dendrite: neurons that respond more to large stimuli receive synaptic inputs on their apical dendrite that cover a larger region of the visual field, are more displaced relative to the receptive field of the neuron, and are more dispersed among each other.

These results suggest a simple interpretation for the effects we had observed after apical pruning. Perhaps in large-preferring neurons the strong responses to large visual stimuli originate from excitatory inputs to the apical dendrite, which cover a large region of the visual field. Losing the apical dendrite, therefore, causes a major loss of input, leading to reduced responses. In small-preferring neurons, instead, the region of the visual field covered by the apical dendrite is small, and perhaps similar in extent to the region covered by other synaptic inputs. In these neurons, the role of the apical dendrite in driving visual responses is redundant with that of other cellular compartments, so that losing this dendrite does not have a major effect.

## Discussion

We showed that in the awake visual cortex, the apical dendrite is critical for spatial integration in neurons that prefer large visual stimuli. In these neurons, pruning the apical dendrite reduces the visual responses, a reduction that is most evident in the responses to large stimuli. Consistently, the apical inputs to large-preferring neurons cover a wider region of visual field than in small-preferring neurons.

Optical pruning provides a precise tool to probe the causal role of individual apical dendrites in the awake brain. Indeed, previous attempts to manipulate the activity of apical dendrites in vivo have targeted entire neural populations ^41,45,45–48^. In doing so, they likely modified not only the activity of the apical dendrites but also the inputs that neurons receive in other compartments. By contrast, the technique of in vivo optical apical pruning ^49^ targets an individual apical dendrite, thus isolating its specific effect on that neuron’s responses.

Applying this technique in the visual cortex revealed a unique visual role for the apical dendrite in cells that integrate signals over a large region. Much interest in visual neuroscience has focused on the phenomenon of size tuning (historically also named hypercomplexity, end-inhibition, surround inhibition, or surround suppression), whereby some neurons respond more to small stimuli than to larger stimuli ^30–34^, while others, particularly in deeper layers prefer larger stimuli ^31,32^. We found that apical pruning substantially reduced responses in large-preferring neurons, suggesting that their apical dendrite provides unique inputs which drive, rather than suppress, the neuron.

Other visual properties, instead, were remarkably immune to apical pruning. While he apical dendrite had been proposed to play a critical role in a neuron’s selectivity for orientation ^60^, Park et al (2019) ^49^ showed that apical pruning largely spares a neuron’s orientation and direction selectivity ^61^. However, in their study, mice were anaesthetized during recordings, potentially hiding the contribution of projections from higher visual areas, whose activity is reduced during sedation ^62,63^. Our data from L5 in the awake cortex (rather than L2/3 under anesthesia) confirms that apical pruning does not affect orientation and direction selectivity. Moreover, apical pruning did not affect the responsiveness and size tuning of neurons that prefer small stimuli.

What mechanisms might explain the resilience of many tuning properties to apical pruning? One possibility is homeostasis. When a cortical neuron detects its activity dropping, it compensates by increasing the strength of its synapses ^64,65^ thus maintaining a constant operating point ^66^. This homeostatic mechanism is likely to engage after apical pruning, strengthening synaptic inputs on the remaining cellular compartments. Further strengthening the effect of existing synaptic inputs is the loss of dendritic surface, which would reduce the neuron’s input conductance ^61^. These phenomena could counteract the effects of apical pruning, but only if the existing inputs are redundant with the lost ones. If the apical tuft receives unique inputs – as appears to be the case for large-preferring neurons – increasing the gain on existing inputs cannot make up for the lost inputs.

Our results suggest that the apical tuft is not a unique site for the suppressive signals that are implicated in size tuning. Size tuning is thought to involve inhibition from SST interneurons, which prefer large stimuli ^34,36,37^ and target the apical tuft^23–27^. Indeed, silencing these cells relieves pyramidal neurons from surround suppression ^37,67^. If the apical inhibition delivered by SST interneurons was the only mechanism supporting size tuning, we should have observed a loss of size tuning in small-preferring neurons, the opposite of what we found. A potential reconciliation of these apparently conflicting results is that size tuning is mostly driven by a reduction in network excitation in the presence of large stimuli ^65^. This reduction may be caused by SST cells, but it would constitute a network effect, not one that could be disrupted by cutting a single apical dendrite.

The results of apical pruning suggest that large-preferring neurons receive on their apical tuft inputs from a larger region of visual space. To test our hypothesis, we combined iGluSnFR imaging of synaptic inputs with calcium imaging of somatic responses to independently measure the visual preference of the soma and of the inputs targeting the apical tuft. Earlier approaches to imaging synaptic activity relied on post-synaptic calcium imaging ^67–70^. Calcium signals in spines, however, can be contaminated by somatic events ^19,68,69^ and largely rely on voltage-gated calcium channel activation and NMDA receptor recruitment ^70–72^, making them less sensitive to weaker inputs, and potentially insensitive to thalamic ones ^73^. Glutamate imaging with iGluSnFR overcomes these limitations ^52,73,74^, revealing the presynaptic activity of a myriad inputs to a target cell.

This technique revealed that large-preferring neurons receive apical inputs that are more broadly distributed, providing an extended source of excitation that correlates with their preference for large stimuli. As expected, apical synaptic inputs were broadly matched to the soma RF location ^75,76^ but locally scattered ^68,77,78^. Crucially, the visual region covered by the inputs to large-preferring cells was significantly larger than for small-preferring cells. The individual inputs, however, had similar receptive field size in the two types of neurons (supplementary Figure 6), suggesting that they might have a common origin.

This distinct spatial organization provides a simple explanation for the vulnerability to the loss of the apical dendrite observed in our causal experiments. Together, these findings demonstrate that the apical dendrite of large-preferring neurons preferentially integrates distal excitatory signals that underlie the neuron’s responses to large stimuli.

We suggest that similar architectures of dendritic connectivity may apply to other cortical areas, governing spatial integration and contextual modulation across modalities. The visual cortex is often used as a model for probing cortical function, thanks in part to the tight correspondence between the visual field and the cortical surface. Similar circuits, however, are thought to be at play throughout the cortex ^79^. If so, our results suggest that in layer 5 of the entire cortex there is a class of pyramidal neurons that integrate the activity of presynaptic neurons scattered over a large and potentially offset cortical region; these distal inputs convey contextual and functionally diverse information preferentially to the apical tuft. In this class of neurons, the apical dendrite serves a fundamental causal role in driving (and not suppressing) the activity of the neurons. Other L5 pyramidal neurons, instead, receive inputs from a more local, and thus functionally homogeneous, cortical region, which do not specifically target the apical tuft. This is one of the many hypotheses that are now testable thanks to the methods of dendritic pruning and glutamate imaging.

## Author contributions

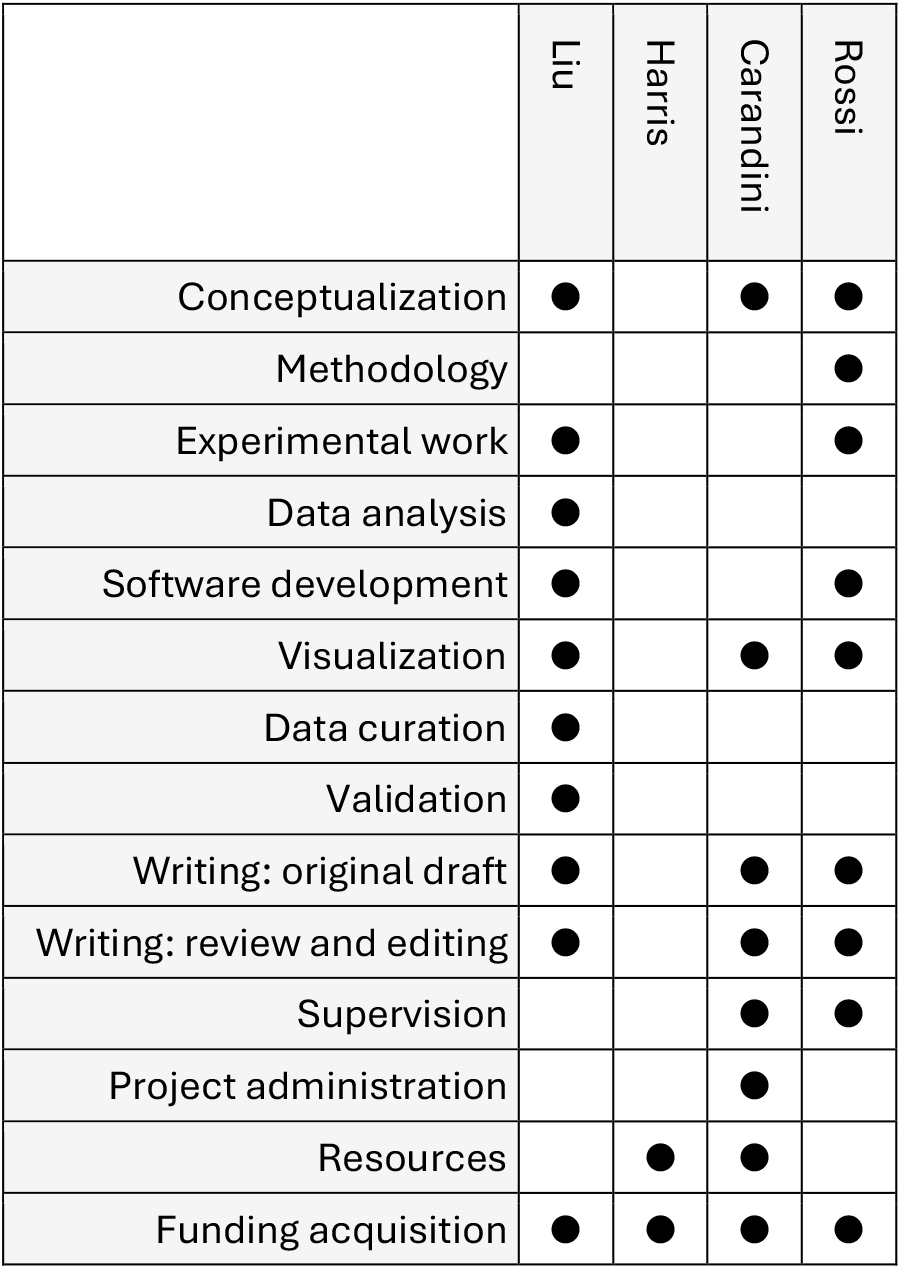

## Acknowledgments

We thank Charu Reddy for laboratory management, Michael Krumin for technical support, George Booth for help with neuronal reconstruction proofreading. This work was supported by grants from Boehringer Ingelheim Fonds (PhD studentship to A.L.), BBSRC, Wellcome Trust (223144/Z/21/Z to M.C. and K.D.H.), UKRI (Frontier Award EP/X022366/1 to M.C.), Armenise-Harvard Foundation (CDA to L.F.R.), and Human Technopole (HT–ECF 3588 to L.F.R.). MC holds the GlaxoSmithKline / Fight for Sight Chair in Visual Neuroscience.

## Methods

All experimental procedures were conducted in accordance with the UK Animals Scientific Procedures Act (1986). Experiments were performed at University College London under personal and project licenses released by the Home Office following appropriate ethics review.

### Mice

Experiments were performed on 21 mice, 7-12 weeks old, of both sexes. The apical pruning experiment was performed on 5 mice from the Rbp4-Cre transgenic line, which expresses Cre in L5 excitatory neurons. Imaging of apical excitatory inputs was performed on 15 wild type (C57bl6/J) mice and one Rbp4-Cre transgenic mouse.

### Surgical procedures

Surgical procedures were carried out under isoflurane anesthesia (1-2% in Oxygen), while the body temperature was monitored and kept at 37-38°C using a closed-loop heating pad, and the eyes were protected with ophthalmic gel (Viscotears Liquid Gel, Alcon Inc.). An analgesic (Rimadyl, 5 mg/kg) was administered subcutaneously on the day of the surgical procedure and on subsequent days, as needed. Dexamethasone (0.5 mg/kg, IM) was administered intramuscularly 30 min prior to the procedure to prevent brain oedema. The head was shaved and disinfected; the cranium was exposed and covered with biocompatible cyanoacrylate glue (Vetbond). First, a stainless-steel head plate with a 10 mm circular opening over the right hemisphere was secured over the skull using dental cement (Super-Bond C&B, 10 Sun Medical Co. Ltd., Japan). Then, a 3mm wide square craniotomy was opened over the right visual cortex (centered at -3.3 mm AP, 2.8 ML from bregma). The exposed brain was irrigated with artificial cerebrospinal fluid at all times (150 mM NaCl, 2.5 mM KCl, 10 mM HEPES, 2 mM CaCl2, 1 mM MgCl2; pH 7.3 adjusted with NaOH, 300 mOsm). Occasionally, a durotomy was performed using a recurved insulin needle, and the dura retracted and removed using forceps. Then, a cocktail of AAV (described in the next section) was injected using a pneumatic injector (Nanoject, Drummond Scientific Company) through a 30-50 µm borosilicate capillary. We typically targeted three 60 nL injections to different V1 sectors, using brain vasculature as a guide. Finally, the craniotomy was sealed with a glass cranial window, attached to the skull using cyanoacrylate glue and dental cement. The window was assembled from a circular cover glass (4 mm diameter, 100 µm thickness) glued to a smaller custom-made insert (3 mm diameter, 300 µm thickness) with index-matched UV curing adhesive (Norland #61).

### Sparse viral expression strategy

In the Rbp-4 Cre mice used in the apical pruning experiments, to co-express GCaMP7s and tdTomato in the same neurons in a sparse ensemble of L5 excitatory neurons, we injected a combination of diluted Cre-dependent FlpO virus (AAV-DIO-FlpO, ∼7 ×10^9^ GC/ml) with high-titer, Flp-dependent reporters AAV-fDIO-GCaMP7s and AAV-fDIO-tdTomato (each ∼1 × 10^12^ GC/ml).

In the wild-type mice used for apical synaptic imaging, to co-express the green iGluSNFRs and the red jRGECO1a in a sparse ensemble of excitatory neurons, we injected at 500 µm cortical depth, a combination of diluted AAV2.1-CaMK2a-Cre (∼10^8^ GC/mL) and concentrated iGluSnFRs variant of one of the three: AAV2.1-hSyn.FLEX.iGluSnFR3.v857.GPI (final concentration: 3.88 ×10^12^ GC /ml; 3 mice); or pGP-AAV-hSyn-flex-iGluSnFR4_v8360-PDGFR-WPRE (2.3 × 10^12^ GC /ml; 1 mouse), or pGP-AAV-hSyn-flex-iGluSnFR4_v8880-PDGFR-WPRE (2.3 × 10^12^ GC /ml; 2 mice). All three were then combined with AAV1.syn.Flex.NES-jRGECO1a.WPRE.SV40 (2.2 × 10^12^ GC /ml).

In one Rbp4-Cre mouse, iGluSnFR4 and jRGECO1a were co-expressed in L5 excitatory neurons restrictedly. This was done by injecting a mixture of diluted AAV-DIO-FlpO (4 × 10^9^ GC/ml) with concentrated Flp-dependent iGluSnFR4 (AAV1-pAAV-hSyn-Flp-FRT-iG-luSnFR4_var8880NGR-WPRE, 4.8 × 10^12^ GC/ml) and jRGECO1a (AAV1-pAAV-hSyn-FlpO-jRGECO1a, 3.8 × 10^12^ GC/ml).

### Two-photon imaging of neuronal responses

Recordings of neuronal activity were performed with a resonant-scanning two-photon microscope (Bergamo II, Thorlabs), equipped with a Nikon 16x, 0.8 NA objective mounted on a piezoelectric z-drive (PIFOC P-725.4CA, Physik Instrumente, range 400 µm) for volumetric multi-plane imaging. The microscope was controlled using ScanImage 2023.1. Excitation light was provided by a femtosecond laser (Chameleon Discovery TPC, Coherent), tuned between 920-1020 nm depending on imaging depth and application. Laser power was depth-adjusted between 30-300 mW according to the piezo position. Sample fluorescence was collected in a green and a red channel: 525/50 nm for the GCaMP6/iGluSnFR3/4, and 605/70 for the tdTomato/jRGECO1a. The imaging objective was light shielded using a custom-made metal cone, a tube, and black cloth to prevent contamination of the fluorescent signal by the visual stimulation light.

During the imaging sessions, mice were head fixed and free to run on a meshed treadmill. For somatic imaging of GCaMP7s, fields of view (FOV) were imaged at 920nm, sampling 512*512 pixels at 30 Hz. Depending on the location of the target sparsely labelled neurons, the FOV spanned on average 265 µm and the objective was scanned across 4-9 imaging planes, separated by 12-40 µm in depth, resulting in an effective sampling rate of 3.3-7.5 Hz. For dendritic imaging of iGluSnFR3 and 4, a single FOV typically spanning 95 µm was imaged at 950 nm, sampling 512*512 pixels at 30 Hz. In both cases, sparsely labelled neurons were used as a reference to target the same cortical volume during this longitudinal imaging.

### Two-photon in vivo morphological reconstructions

For morphological reconstructions of the full dendritic tree, structural z-stacks were acquired by scanning the piezoelectric z-drive attached to the objective in steps of 1 µm, thence imaging 100-200 repetitions of the same cortical volume. Z-stacks typically spanned a FOV of 600*600 µm in apical pruning experiments, and a FOV of 120*120 µm in glutamate imaging experiments, imaged with a lateral resolution of greater or equal to 1 pixels/µm. Repetitions were motion corrected and averaged to obtain a high SNR imaging volume: we first registered and averaged repetitions of individual imaging planes using an FFT-based subpixel registration algorithm ^80^; then we registered consecutive planes to ensure vertical alignment; finally, to correct for bleed-through of GCaMP7 fluorescence into the red channel, a scaled version of the green channel was subtracted, with the scaling factor estimated via robust regression using the brightest 20% of pixels ^81^. Due to the limited range of the piezoelectric z-drive (300-400 µm), two adjacent z-stacks were acquired and subsequently concatenated across depth reconstruct both the apical dendrites and somas. Before concatenation, stacks were normalized to a common intensity range, and aligned laterally using subpixel phase correlation ^82^.

We used volumetric images to reconstruct neuronal morphologies with NeuTube ^55^; we then imported and further analyzed reconstructions in MATLAB with the Trees toolbox ^83^.

### Two-photon dendritic pruning

A neuron was considered for the dendrite pruning experiment when both its visual tuning and its morphology met two criteria. The first was clear orientation or direction selectivity (OSI or DSI >0.2) and size preference (P<0.05 in oneway ANOVA test across sizes). The second is based on the visibility of the apical dendrite in the morphological reconstruction and during the in vivo imaging. Specifically, its ascending apical trunk needed to be clearly visible when excited at 950 nm at modest laser power. All neurons passed such selection were randomly assigned with a label of ‘prune’ or ‘control’ with equal chance. Control neurons shared the same cortical space as the ones to be pruned, while their apical dendrite would not be pruned.

Targeted optical dendritic pruning ^49^ was performed under the same microscope used for imaging, tuned at 800 nm. To optically cut the apical trunk, we centered the target at high magnification and fixed the microscope mirrors to point a stationary beam in the middle of the FOV. A custom the MATLAB script automatically switched off the PMTs, enabled the laser via the AOM to deliver pulses of high intensity illumination at 800 nm (duration: 5-10s; power: 40-150 mW) while ensuring the PMTs are switched off. The system was then immediately reverted to resonant frame-scanning mode to assess the pruning outcome. Successful photoablation was assessed by imaging at 920 nm: it resulted in elevated calcium in the soma, loss of structural continuity in the trunk, and progressive swelling of the dendritic stub. Successful photoablation was confirmed with a structural z-stack on the day following the procedure. If the unsuccessful, the procedure was repeated with small power increments, limiting cumulative exposure to five attempts to prevent collateral damage on the surrounding tissue. Final confirmation of the lesion was performed via structural z-stacks acquired the following day.

### Visual stimulation

Visual stimuli were generated in MATLAB (MathWorks) using the Psychophysics Toolbox^84^ and displayed on 3 gamma-corrected LCD monitors (Adafruit 1.8” TFT LCD Display, resolution 128×160 px) arranged at 90^°^ to each other. The screens were covered with Fresnel lenses to correct for viewing angle inhomogeneity of the LCD intensity. The mouse was positioned at the center of the layout, so that the monitors spanned ±135 deg horizontally and ±35 deg vertically. Stimuli were only presented in the left hemifield, contralateral to the right hemisphere.

To map the retinotopy of the imaged area and estimate the RF of neurons, we presented sparse, spatial white noise stimuli, consisting of patterns of black and white squares (6 deg x 6 deg each) on a gray background, refreshing at 5 Hz, typically in 10 min sequences repeated 3 times for each recording. At any point in time, each square had a 2% probability of being nongray, independent of the other squares.

To measure direction and size tuning, we presented drifting gratings of different directions and diameters (2 s duration; 100% contrast; spatial frequency 0.05 cycles/deg; temporal frequency 1 Hz), centered at the centroid of all neurons’ RFs in each FOV. Two stimulus sets were used. Set 1 comprised 64 combinations of 8 directions (0–315°, 45° steps) and 8 sizes (5–60 deg), plus blank trials. Set 2 comprised 32 combinations of the same 8 directions and 4 sizes (5, 20, 40, 80 deg), plus blanks. Blank trials were treated as a stimulus with 0 deg in size. Stimuli were presented in pseudorandom order with an inter-stimulus interval of 2–3 s. Each condition was repeated 8 times in Set 1 and 20 times in Set 2. For a given field of view, only one stimulus set was used, and the stimulus order was randomized across sessions.

### Processing of two-photon data

Two-photon imaging data were processed using Suite2p ^85^. For somatic GCaMP7s recordings, the pipeline included rigid image registration, segmentation of active region of interest (ROIs) at the scale of neuronal somas, and estimation of the neuropil signal contaminating each ROI. The final selection of ROIs was curated manually to select target neurons expressing tdTomato, identified by inspecting the co-registered average tdTomato image, and discard spurious or noisy ROIs.

The neuropil signal was weighted by a correction factor α, determined separately for each ROI ^86^, before being subtracted. The correction factor was estimated from the linear relationship between the lowest somatic fluorescence compatible with any value of fluorescence in the neuropil. For each neuron i the neuropil signal Ni(t) was binned into 20 intervals; for each interval, the 5th percentile of the matching time-points of raw somatic fluorescence Fi(t) was measured as an estimate of baseline fluorescence. The correction factor αi was then computed by linear regression between the median of each neuropil interval and estimates of baseline fluorescence, to fit the lower envelope of the scatterplot of neuropil versus somatic fluorescence. The corrected fluorescence was computed as Fi(t) – αiNi(t).

To compare activity across longitudinal recordings, somatic responses were normalized as the relative change from baseline fluorescence (dF/F0). F0 was calculated as the intercept of relationship between neuropil and raw somatic fluorescence. This normalization accounted for potential difference in imaging parameters across days.

For iGluSnFR3/4 recordings, we used instead two-steps non-rigid registration, and segmentation of active ROIs at the spatial scale expected from synapses. Neuropil correction was not necessary for sparse synaptic imaging, and responses were z-scored before further analysis.

### Analysis of direction, orientation and size tuning

Stimulus triggered responses time-courses were computed as the difference between the fluorescent trace and the average baseline activity 2 s prior to stimulus presentation, then averaged across trials of stimuli of the same type. Responses were then quantified as the average fluorescence in a 4 s window following stimulus, and averaged across stimuli of the same directions and blank-subtracted to obtain direction tuning curves, or of the same size to obtain size tuning curves. The preferred orientation, direction and size were defined as those eliciting the maximal response. In sessions where we presented 8 rather than 4 sizes (n=12), responses were interpolated to match the rest of the data.

Direction tuning curves were parametrized as double Gaussian distributions and fit using least-square optimization, and the resulting best-fit tuning curves were evaluated at 5° angular resolution. Tuning width was quantified as the half-width at half-maximum of the minimum-subtracted tuning function.

To estimate the center of mass (CoM) for size tuning curves responses across stimulus sizes (n = 5, including a size 0 blank control) were interpolated onto a fine 2,000-point linear grid spanning 0 deg to the maximum tested size using piecewise cubic Hermite interpolating polynomials (*pchip*, Matlab). Negative responses were clamped to zero. The spatial CoM was calculated by summing the position-weighted responses across the grid and dividing by the total area under the curve:

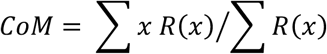

where x represents the stimulus size and R(x) is the interpolated, non-negative response.

### Permutation test for the effects of pruning on visual responses

Visual responses from the ‘pre’ and the ‘post’ recording sessions were normalized to the maximum response recorded in the ‘pre’ session. For response amplitudes, the session-wise change was quantified as the ratio between ‘pre’ and ‘post’ response maxima; for tuning width, the session-wise change was quantified as the difference between ‘post’ and ‘pre’ widths. To assess the effect of pruning in different groups of neurons (pruned vs control, or small-preferring vs large-preferring neurons), we used a group label-permutation test. The observed difference in group mean was evaluated against a null distribution generated by randomly shuffling group labels (10,000 iterations) and recomputing the group difference each time. P-values were calculated as the fraction of shuffled differences that exceeded the observed value in the hypothesized direction (pruned – control giving more negative metrics).

### Receptive field analysis

Receptive fields (RFs) were estimated using regularized linear regression between neural responses (z-scored) and the stimulus history ^87^. The stimulus design matrix incorporated time-lagged predictors (0–1.4 s) using a Toeplitz structure. Smoothness of the RF was enforced by penalizing the Laplacian of the filter, with the regularization parameter (λ) selected via 3-fold cross-validation to maximize predictive performance.

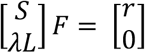

Model performance was quantified using explained variance (EV), and final RFs were fit using the optimal λ on the full dataset. RF significance was assessed using a circular shift test, in which stimulus predictors were temporally shuffled (1,000 iterations) to generate a null distribution; p-values were computed as the fraction of null EVs exceeding the observed EV.

In the apical input imaging experiment, we eliminated noisy synaptic ROIs with ill-defined RFs, by applying several criteria. For iGluSnFR4, ROIs were classified as responsive if EV > 0.15 and p < 0.005. For iGluSnFR3, the criteria were EV > 0.05 and p < 0.01. Spatial RFs were defined as the response amplitudes at the time slice where maximal response occurred, interpolated at 1deg resolution, and then fit with a 2D Gaussian.

To characterize the spatial organization and distribution of synaptic RFs for each neuron, while reducing the influence of outliers, we computed the robust multivariate location and covariance matrix. For neurons with enough synaptic RFs (n>15), this estimation was performed using the Minimum Covariance Determinant method via MATLAB’s *robustcov* function. For neurons with low synaptic RF counts (n<15), where robust covariance estimation is underdetermined, the centroid was defined by the spatial median coordinate, and the covariance matrix was calculated using standard sample covariance. In both cases, a minor diagonal regularization term proportional to the trace of covariance matrix was added to ensure numerical stability and positive definiteness. The total area of this synaptic footprint was calculated as *πab*, where half of the major axis length (*a*) and of the minor axis (*b*) were determined directly from the covariance matrix eigenvalues scaled to a 90% confidence threshold.

Apical synaptic and somatic RF offsets were quantified on a per-neuron basis by computing 2D vectors from the somatic RF center to each synaptic RF. The mean offset vector (component-wise average) was used to summarize the population, with its magnitude reflecting the average displacement of synaptic RFs relative to the soma in visual space.

To evaluate whether the offset between somatic and apical compound RFs could arise from differences in spatial statistics or count of synaptic RF, we implemented a zero-centered Monte Carlo population null model. For each neuron, we generated synthetic synaptic populations matching the neuron’s empirical synaptic count from multivariate distributions with the empirical spatial covariance, but centered at the somatic RF. Across 10,000 iterations, simulated offset magnitudes were averaged across neurons to establish a population null distribution of expected group mean differences. The empirical group difference was then compared against this null distribution. Twotailed Monte Carlo p-values were calculated as the proportion of simulated iterations where the absolute null group difference equaled or exceeded the absolute observed difference.

## Supplementary Figures

**Supplementary Figure 1.**
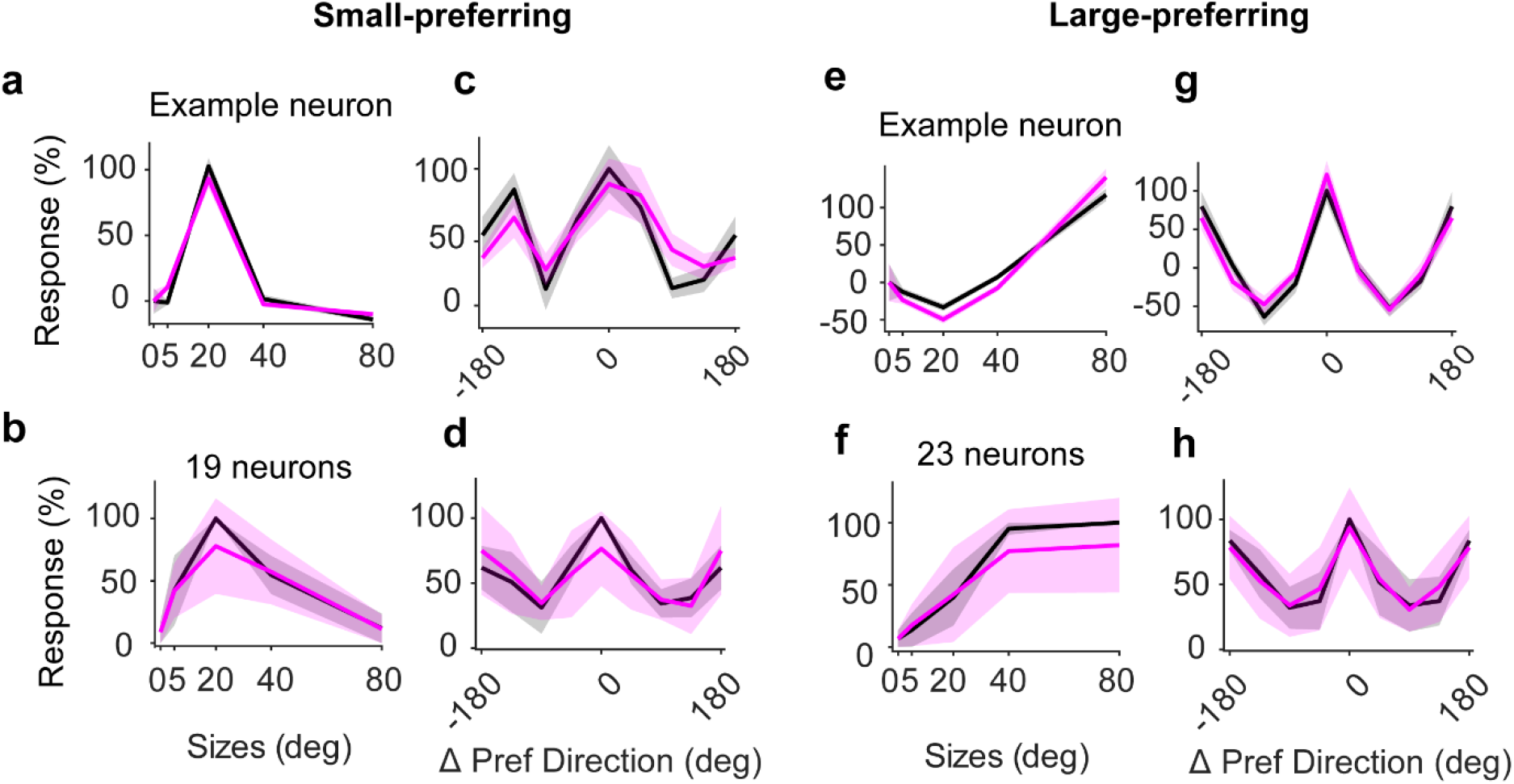
Control neurons maintain stable visual tuning after pruning neighboring dendrites. (**a**) Tuning for stimulus size of an example small-preferring neuron (neuron 17), showing the mean responses before (black) and after (magenta) pruning neighboring dendrites. At each size, the responses are averaged over all stimulus orientations and directions. Responses are normalized so that 0 is the response to a blank screen (measured separately in Pre and Post sessions) and 100% is the response to the best stimulus (measured in the Pre session). Shaded regions show ± s.e. (standard error) across repeats. (**b**) Same, for the population of pruned small-preferring neurons (giving strongest responses to 5 deg and 20 deg stimuli, n=19). Curves show median values, and shaded areas show ± 1 m.a.d. (median absolute deviation) across neurons. (**c**) Same as **a**, showing the example neuron’s tuning for stimulus orientation and direction. At each orientation and direction, the responses are averaged over all sizes. (**d**) Same as (b), showing the population’s tuning for stimulus orientation and direction. (**e-h**) Same as **a-f**, for neurons preferring large stimuli (40 or 80 deg), showing an example neuron (**g, j**) and the population of control neurons (n = 23, **f, h**).

**Supplementary Figure 2.**
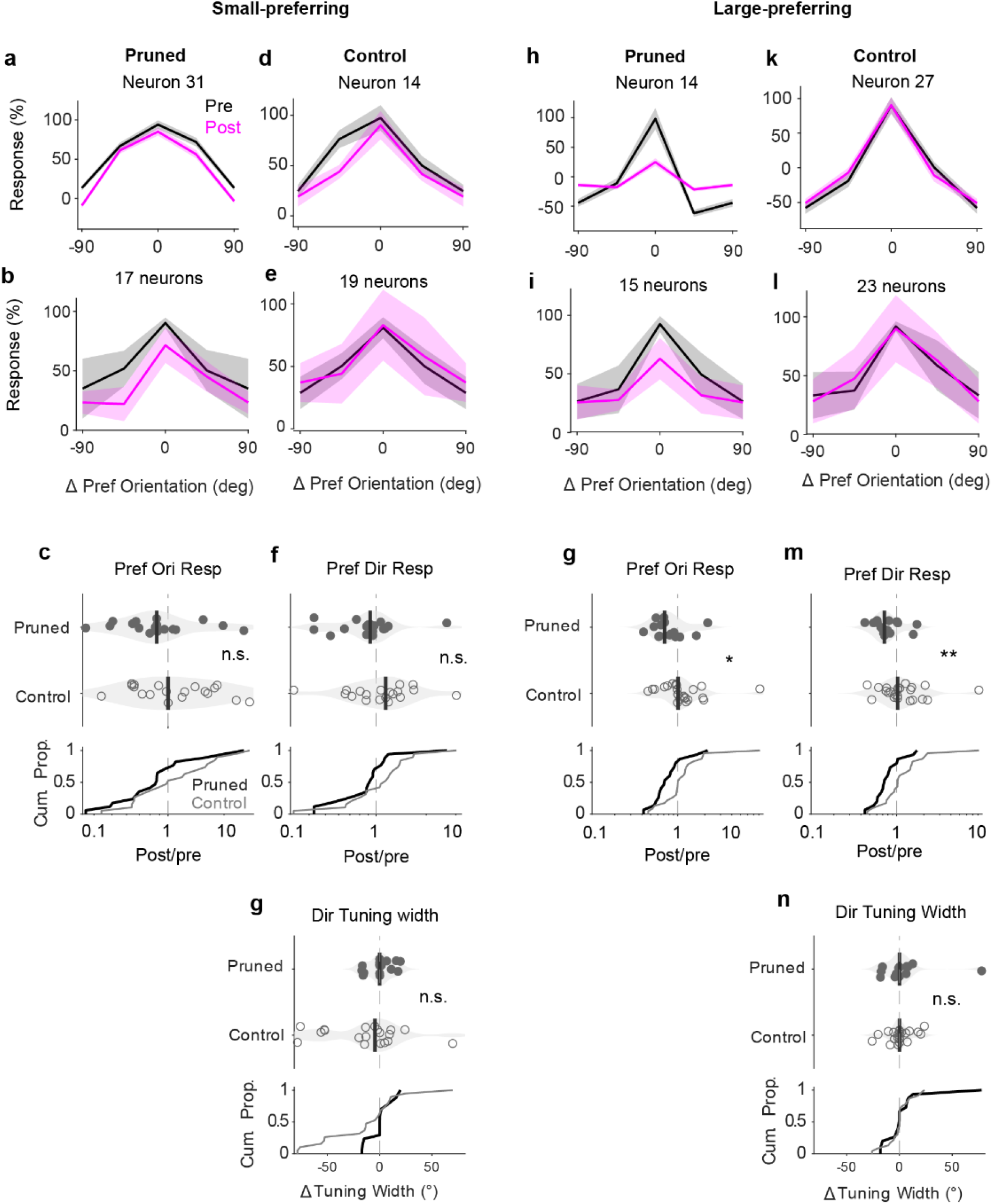
Pruning reduces responses to preferred orientation and direction in large-preferring neurons, while sparing orientation and direction selectivity. (**a**) Tuning for stimulus orientation of an example pruned small-preferring neuron (neuron 31), showing the mean responses before (black) and after (magenta) apical pruning. (**b)** Same, for the population of pruned small-preferring neurons (n=17). (**c**) Effect of pruning on responses at the preferred orientation, for pruned small-preferring neurons (n=17, filled dots) and control small-preferring neurons (n=19, hollow dots). (Median effect: control vs pruned small-preferring: 1.00 vs 0.870, p = 0.097, (n.s.) Permutation test). (**d**-**e**) Same as a and b, for an example (**d**) and the population (**e**) of small-preferring (n=19) control neurons. (**f**) Same as c, for the effect on responses at the preferred direction. (Median effect: control vs pruned small-preferring: 1.14 vs 0.924, p = 0.132, (n.s.) Permutation test). (**g**) Same as **c**, showing the effect of pruning on direction tuning width. (Median effect: control vs pruned small-preferring: -4.61 vs -0.00175, p = 0.844, (n.s.) Permutation test). (**h**-**n**) Same as (**a**-**f**) for large-preferring neurons, showing two example neurons (**h, k**) and the population of pruned (n=15, **i**) and control neurons (n = 23, **l**). (**g**: effect on responses at preferred orientation, median effect: control vs pruned large-preferring: 1.02 vs 0.704, p = 0.025, (*) Permutation test; **m**: effect on responses at preferred direction control vs pruned large-preferring: 0.034 vs -0.109, p = 0.009, (**) Permutation test; **n**: effect on direction tuning width control vs pruned large-preferring: - 0.00172 vs -1.35e-06 p = 0.725, (n.s.) Permutation test).

**Supplementary Figure 3.**
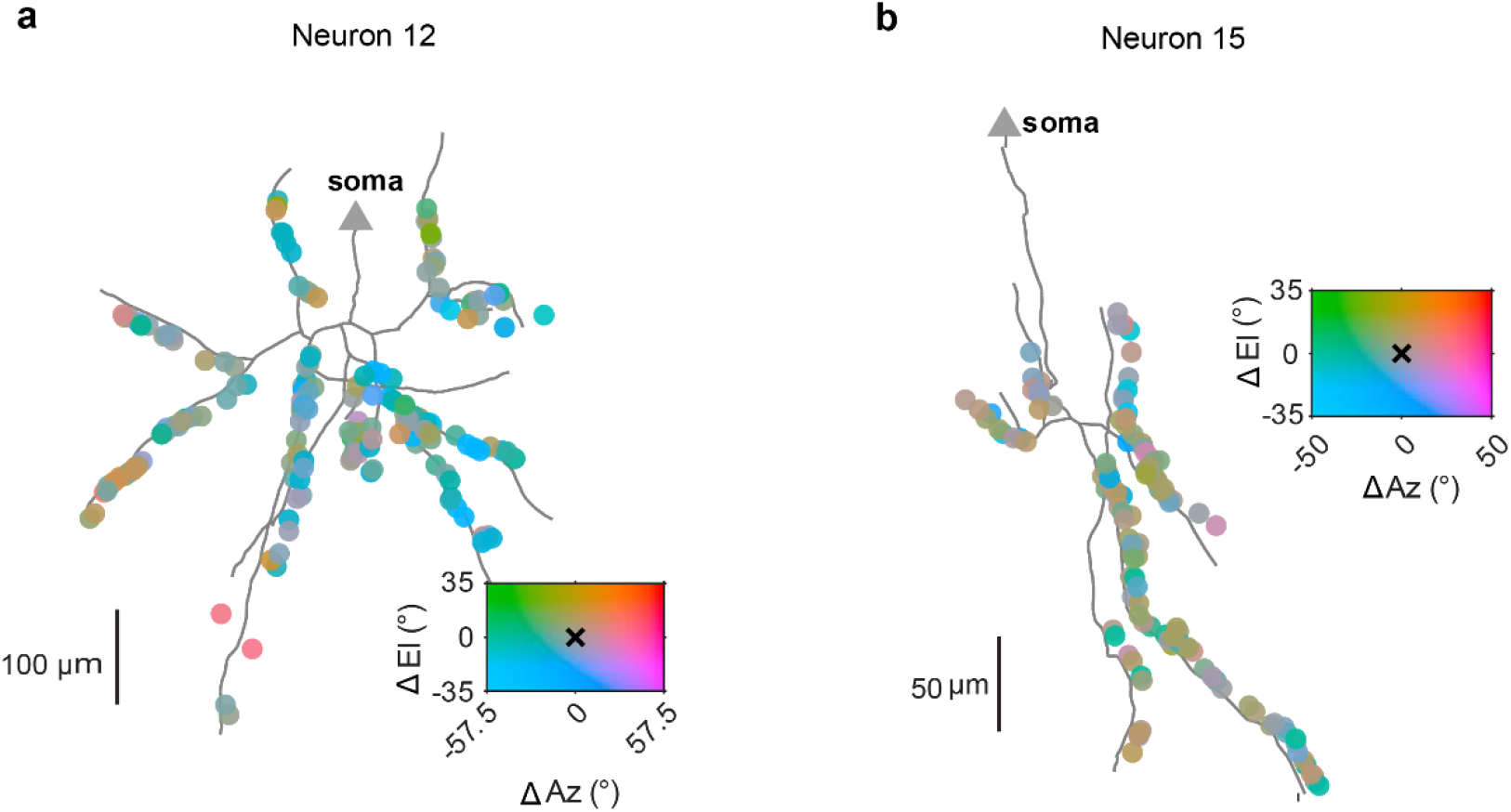
Local heterogeneity in apical inputs. (**a**) Spatial map of apical tuft inputs recorded from a single neuron (small-preferring), stitched across sessions. Inputs (dots) are color coded according to receptive field distance from the soma (triangle), overlaid on the morphological reconstruction of the apical dendritic tree obtained from a structural z-stack. Scale bar: anatomical distance in cortical space (100 µm). RF distance in azimuth and elevation is mapped with a 2D colormap centered on the somatic RF (crosshair). (**b**) Same as **a** for a second example neuron (small-preferring).

**Supplementary Figure 4.**
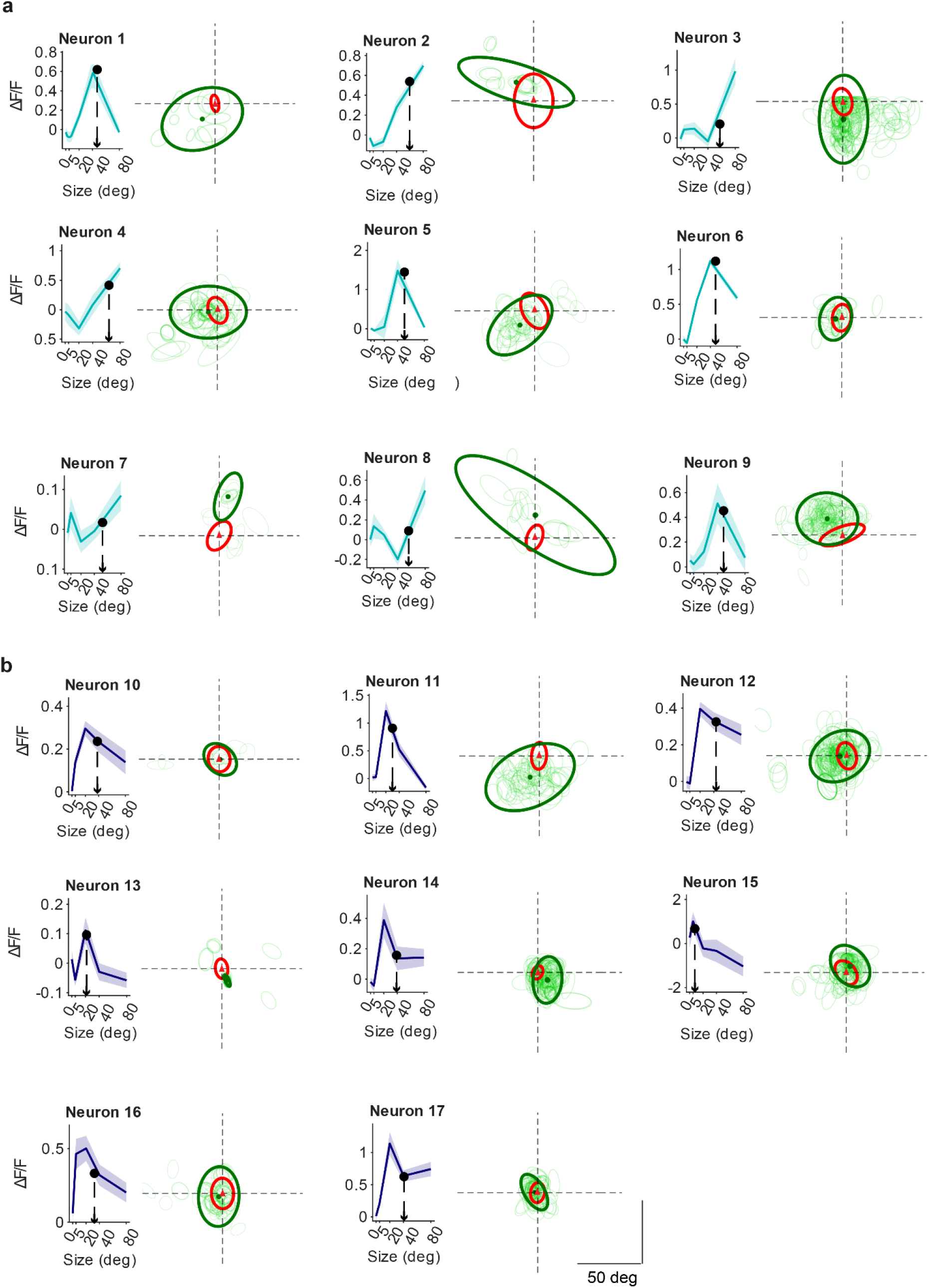
Somatic and apical RFs from all imaged neurons. **(a)** Size tuning (left) and RF data (right) for each large preferring neuron (n = 9, teal). In the somatic size tuning curve, the shaded area indicates the standard error across stimulus repeats; the center of mass of the responses (arrow) constitutes the mean responsive size. RF data reports the apical RFs (light green), the compound apical RF (dark green ellipse, centered on green dot) relative to the center of the somatic RF (red ellipse centered on the origin). (**b**) Same as **a**, for small-preferring neurons (n = 8, purple).

**Supplementary Figure 5.**
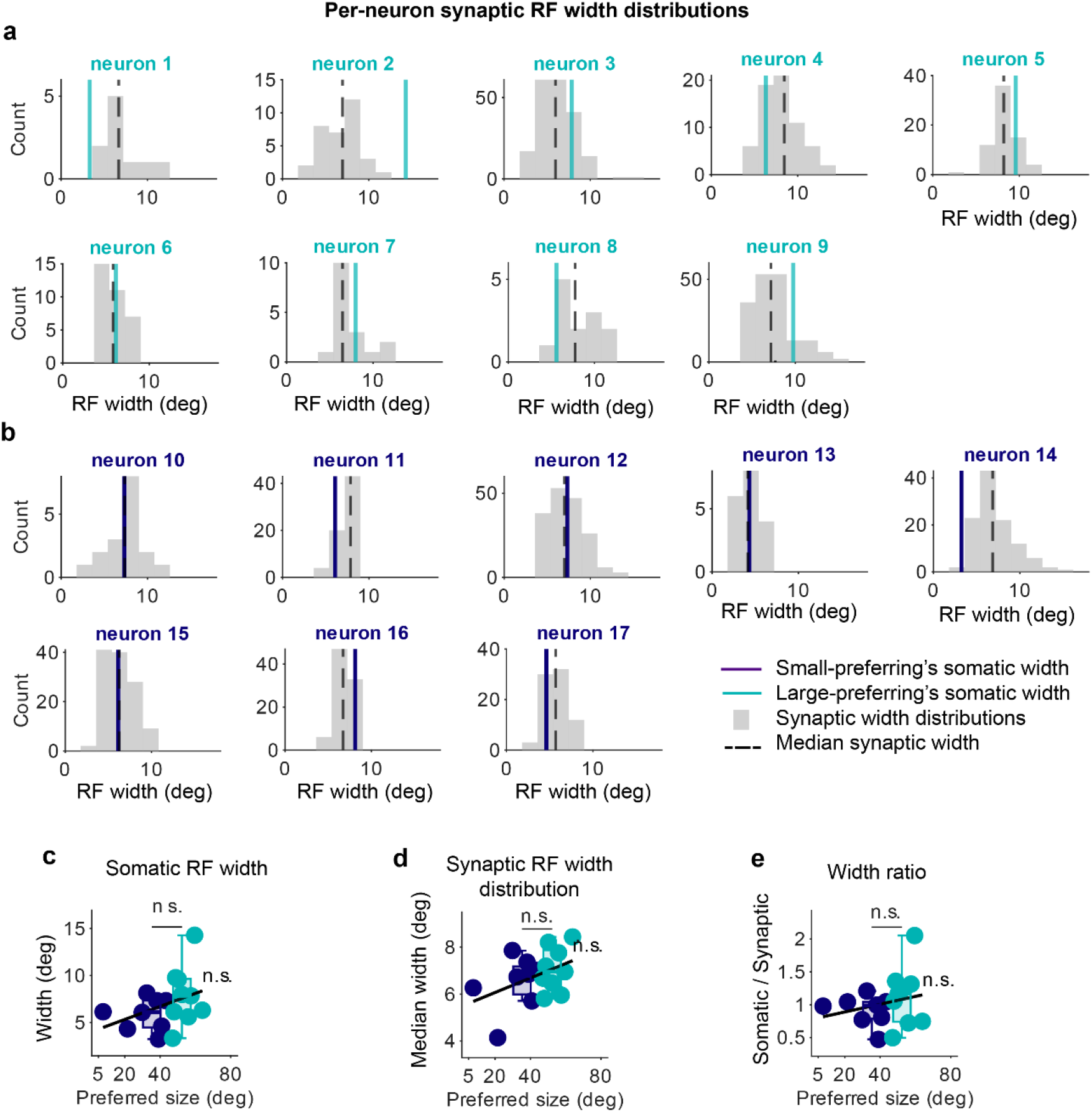
Apical and somatic RF extent do not scale with somatic preferred grating size. (**a**) Comparison of somatic and apical receptive fields (RFs) extent for all recorded large-preferring neurons. Histograms (gray) show the distribution RF extent of apical synaptic inputs. RF extent is measured as the mean of the horizontal and vertical axis of the ellipse fit to the RF spatial footprint. The solid-colored line indicates the somatic RF extent, and the dashed black line denotes the median apical RF extent. (**b**) Same for small-preferring neurons and their apical inputs. **(b)** The size of somatic RF extent vs mean somatic responsive size for small-preferring (purple) and large-preferring (teal) neurons. Each dot represents a neuron, overlaid with group boxplots. Lines indicate linear regression across all neurons. We found no significant linear relation or between-group differences (Wilcoxon rank-sum test; n.s., not significant). (**d**) Same, for the median extent of apical RFs vs mean somatic responsive size. (**e**) Same for the ratio between somatic and median apical RF extent vs mean somatic responsive size.

**Supplementary Figure 6.**
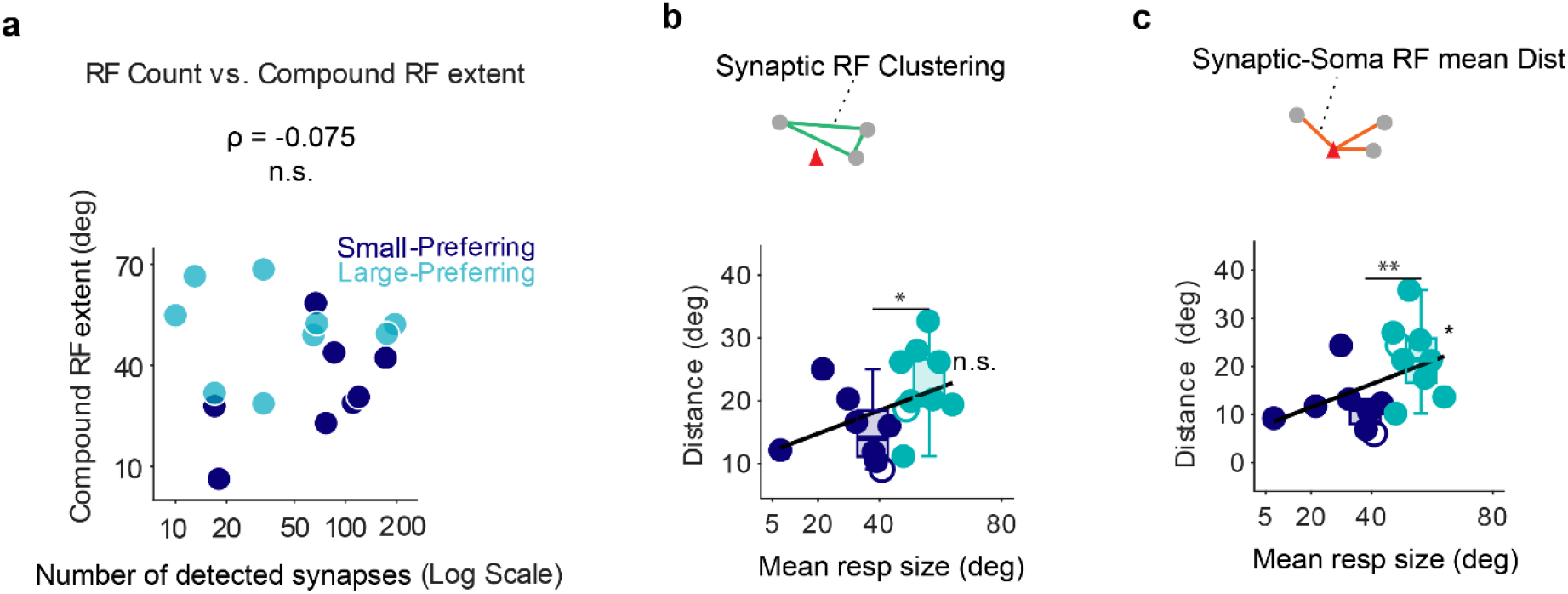
Apical RF organization, mean response size and number of sampled synapses. (**a**) The total number of detected apical RFs (log scale) vs the extent of compound RF (degrees) for for small-preferring (purple) and large-preferring (teal) neurons. Difference in extent of the compound apical RF between large- and small-preferring neurons does not depend on the number of sampled apical inputs (2-ways ANOVA, p=0.030 for group, p= 0.60 for number of synapse count; Spearman’s rank correlation: ρ = -0.075, p = 0.78 (n.s.)). **(b)** Pairwise distance between synaptic RFs (RF clustering) vs mean responsive size for small-preferring (purple) and large-preferring (teal) neurons. Each dot represents a neuron (hollow dots: example neuron from Figure 4 **a, b**), overlaid with group boxplots. Lines indicate linear regression across all neurons. Asterisks indicate slope significance and between-group differences (Wilcoxon ranksum test; *p < 0.05, **p < 0.01, n.s., not significant). **(c)** Same, for the mean distance between the somatic RF and all the synaptic RFs.

## References

1. Spruston, N. Pyramidal neurons: dendritic structure and synaptic integration. Nat Rev Neurosci 9, 206–221 (2008).

2. Ledderose, J. M. T. et al. Layer 1 of somatosensory cortex: an important site for input to a tiny cortical compartment. Cereb Cortex 33, 11354–11372 (2023).

3. Rubio-Garrido, P., Pérez-de-Manzo, F., Porrero, C., Galazo, M. J. & Clascá, F. Thalamic input to distal apical dendrites in neocortical layer 1 is massive and highly convergent. Cereb Cortex 19, 2380–2395 (2009).

4. Cauller, L. Layer I of primary sensory neocortex: where top-down converges upon bottom-up. Behav Brain Res 71, 163–170 (1995).

5. Felleman, D. J. & Van Essen, D. C. Distributed hierarchical processing in the primate cerebral cortex. Cereb Cortex 1, 1–47 (1991).

6. Petreanu, L., Mao, T., Sternson, S. M. & Svoboda, K. The subcellular organization of neocortical excitatory connections. Nature 457, 1142–1145 (2009).

7. Rockland, K. S. & Pandya, D. N. Laminar origins and terminations of cortical connections of the occipital lobe in the rhesus monkey. Brain Research 179, 3–20 (1979).

8. Schuman, B., Dellal, S., Prönneke, A., Machold, R. & Rudy, B. Neocortical Layer 1: An Elegant Solution to Top-Down and Bottom-Up Integration. Annu Rev Neurosci 44, 221–252 (2021).

9. Larkum, M. A cellular mechanism for cortical associations: an organizing principle for the cerebral cortex. Trends in Neurosciences 36, 141–151 (2013).

10. Takahashi, N. et al. Active dendritic currents gate descending cortical outputs in perception. Nat Neurosci 23, 1277–1285 (2020).

11. Xu, N. et al. Nonlinear dendritic integration of sensory and motor input during an active sensing task. Nature 492, 247–251 (2012).

12. Major, G., Larkum, M. E. & Schiller, J. Active Properties of Neocortical Pyramidal Neuron Dendrites. Annu. Rev. Neurosci. 36, 1–24 (2013).

13. Amitai, Y., Friedman, A., Connors, B. W. & Gutnick, M. J. Regenerative Activity in Apical Dendrites of Pyramidal Cells in Neocortex. Cereb Cortex 3, 26–38 (1993).

14. Grienberger, C., Chen, X. & Konnerth, A. NMDA Receptor-Dependent Multidendrite Ca 2+ Spikes Required for Hippocampal Burst Firing In Vivo. Neuron 81, 1274–1281 (2014).

15. Larkum, M. E. Top-down Dendritic Input Increases the Gain of Layer 5 Pyramidal Neurons. Cerebral Cortex 14, 1059–1070 (2004).

16. Larkum, M. E., Nevian, T., Sandler, M., Polsky, A. & Schiller, J. Synaptic Integration in Tuft Dendrites of Layer 5 Pyramidal Neurons: A New Unifying Principle. Science 325, 756–760 (2009).

17. Larkum, M. E., Zhu, J. J. & Sakmann, B. Dendritic mechanisms underlying the coupling of the dendritic with the axonal action potential initiation zone of adult rat layer 5 pyramidal neurons. The Journal of Physiology 533, 447–466 (2001).

18. Schiller, J., Schiller, Y., Stuart, G. & Sakmann, B. Calcium action potentials restricted to distal apical dendrites of rat neocortical pyramidal neurons. J Physiol 505 (Pt 3), 605–616 (1997).

19. Yuste, R., Gutnick, M. J., Saar, D., Delaney, K. R. & Tank, D. W. Ca2+ accumulations in dendrites of neocortical pyramidal neurons: An apical band and evidence for two functional compartments. Neuron 13, 23–43 (1994).

20. Abs, E. et al. Learning-Related Plasticity in Dendrite-Targeting Layer 1 Interneurons. Neuron 100, 684–699.e6 (2018).

21. Hestrin, S. & Armstrong, W. E. Morphology and physiology of cortical neurons in layer I. J Neurosci 16, 5290–5300 (1996).

22. Palmer, L., Murayama, M. & Larkum, M. Inhibitory Regulation of Dendritic Activity in vivo. Front. Neural Circuits 6, (2012).

23. Fino, E. & Yuste, R. Dense inhibitory connectivity in neocortex. Neuron 69, 1188–1203 (2011).

24. Kawaguchi, Y. GABAergic cell subtypes and their synaptic connections in rat frontal cortex. Cerebral Cortex 7, 476–486 (1997).

25. Silberberg, G. & Markram, H. Disynaptic Inhibition between Neocortical Pyramidal Cells Mediated by Martinotti Cells. Neuron 53, 735–746 (2007).

26. Tremblay, R., Lee, S. & Rudy, B. GABAergic Interneurons in the Neocortex: From Cellular Properties to Circuits. Neuron 91, 260–292 (2016).

27. Wang, Y. et al. Anatomical, physiological and molecular properties of Martinotti cells in the somatosensory cortex of the juvenile rat. J Physiol 561, 65–90 (2004).

28. Kapfer, C., Glickfeld, L. L., Atallah, B. V. & Scanziani, M. Supralinear increase of recurrent inhibition during sparse activity in the somatosensory cortex. Nat Neurosci 10, 743–753 (2007).

29. Pfeffer, C. K., Xue, M., He, M., Huang, Z. J. & Scanziani, M. Inhibition of inhibition in visual cortex: the logic of connections between molecularly distinct interneurons. Nat Neurosci 16, 1068–1076 (2013).

30. Angelucci, A. et al. Circuits and Mechanisms for Surround Modulation in Visual Cortex. Annu Rev Neurosci 40, 425–451 (2017).

31. Cavanaugh, J. R., Bair, W. & Movshon, J. A. Selectivity and Spatial Distribution of Signals From the Receptive Field Surround in Macaque V1 Neurons. Journal of Neuro-physiology 88, 2547–2556 (2002).

32. Gilbert, C. D. Laminar differences in receptive field properties of cells in cat primary visual cortex. The Journal of Physiology 268, 391–421 (1977).

33. Hubel, D. H. & Wiesel, T. N. Receptive fields and functional architecture in two nonstriate visual areas (18 and 19) of the cat. Journal of Neurophysiology 28, 229–289 (1965).

34. Niell, C. M. & Scanziani, M. How Cortical Circuits Implement Cortical Computations: Mouse Visual Cortex as a Model. Annu Rev Neurosci 44, 517–546 (2021).

35. Ozeki, H. et al. Relationship between Excitation and Inhibition Underlying Size Tuning and Contextual Response Modulation in the Cat Primary Visual Cortex. J. Neurosci. 24, 1428–1438 (2004).

36. Adesnik, H., Bruns, W., Taniguchi, H., Huang, Z. J. & Scanziani, M. A neural circuit for spatial summation in visual cortex. Nature 490, 226–231 (2012).

37. Hendricks, W. D., Sadahiro, M., Mossing, D., Veit, J. & Adesnik, H. Feature-tuned synaptic inputs to somatostatin interneurons drive context-dependent processing. Neuron 114, 1257–1268.e5 (2026).

38. Pérez-Garci, E., Larkum, M. E. & Nevian, T. Inhibition of dendritic Ca2+ spikes by GABAB receptors in cortical pyramidal neurons is mediated by a direct Gi/o-β-subunit interaction with Cav1 channels. J Physiol 591, 1599–1612 (2013).

39. Benezra, S. E., Patel, K. B., Perez Campos, C., Hillman, E. M. & Bruno, R. M. Learning enhances behaviorally relevant representations in apical dendrites. eLife 13, (2024).

40. Fişek, M. et al. Cortico-cortical feedback engages active dendrites in visual cortex. Nature 617, 769–776 (2023).

41. Manita, S. et al. A Top-Down Cortical Circuit for Accurate Sensory Perception. Neuron 86, 1304–1316 (2015).

42. Beaulieu-Laroche, L., Toloza, E. H. S., Brown, N. J. & Harnett, M. T. Widespread and Highly Correlated Somato-dendritic Activity in Cortical Layer 5 Neurons. Neuron 103, 235–241.e4 (2019).

43. Francioni, V., Padamsey, Z. & Rochefort, N. L. High and asymmetric somato-dendritic coupling of V1 layer 5 neurons independent of visual stimulation and locomotion. eLife 8, e49145 (2019).

44. Otor, Y. et al. Dynamic compartmental computations in tuft dendrites of layer 5 neurons during motor behavior. Science 376, 267–275 (2022).

45. Doron, G. et al. Perirhinal input to neocortical layer 1 controls learning. Science 370, eaaz3136 (2020).

46. Maristany De Las Casas, E. et al. Tuft dendrites in frontal motor cortex enable flexible learning. Science 392, eadx4358 (2026).

47. Schoenfeld, G. et al. Unsigned temporal difference errors in cortical L5 dendrites during learning. bioRxiv (2021).

48. Takahashi, N., Oertner, T. G., Hegemann, P. & Larkum, M. E. Active cortical dendrites modulate perception. Science 354, 1587–1590 (2016).

49. Park, J. et al. Contribution of apical and basal dendrites to orientation encoding in mouse V1 L2/3 pyramidal neurons. Nat Commun 10, 5372 (2019).

50. Harris, K. D. & Mrsic-Flogel, T. D. Cortical connectivity and sensory coding. Nature 503, 51–58 (2013).

51. Aggarwal, A. et al. Glutamate indicators with increased sensitivity and tailored deactivation rates. Nat Methods 23, 417–425 (2026).

52. Gerfen, C. R., Paletzki, R. & Heintz, N. GENSAT BAC cre-recombinase driver lines to study the functional organization of cerebral cortical and basal ganglia circuits. Neuron 80, 1368–1383 (2013).

53. Dana, H. et al. High-performance calcium sensors for imaging activity in neuronal populations and microcompartments. Nat Methods 16, 649–657 (2019).

54. Feng, L., Zhao, T. & Kim, J. neuTube 1.0: A New Design for Efficient Neuron Reconstruction Software Based on the SWC Format. eneuro 2, ENEURO.0049-14.2014 (2015).

55. Siegle, J. H. et al. Survey of spiking in the mouse visual system reveals functional hierarchy. Nature 592, 86–92 (2021).

56. Murgas, K. A., Wilson, A. M., Michael, V. & Glickfeld, L. L. Unique Spatial Integration in Mouse Primary Visual Cortex and Higher Visual Areas. J. Neurosci. 40, 1862–1873 (2020).

57. Yeh, C.-I., Xing, D., Williams, P. E. & Shapley, R. M. Stimulus ensemble and cortical layer determine V1 spatial receptive fields. Proc Natl Acad Sci U S A 106, 14652–14657 (2009).

58. Cavanaugh, J. R., Bair, W. & Movshon, J. A. Nature and Interaction of Signals From the Receptive Field Center and Surround in Macaque V1 Neurons. Journal of Neurophysiology 88, 2530–2546 (2002).

59. Smith, S. L., Smith, I. T., Branco, T. & Häusser, M. Dendritic spikes enhance stimulus selectivity in cortical neurons in vivo. Nature 503, 115–120 (2013).

60. Petousakis, K.-E., Park, J., Papoutsi, A., Smirnakis, S. & Poirazi, P. Modeling apical and basal tree contribution to orientation selectivity in a mouse primary visual cortex layer 2/3 pyramidal cell. eLife 12, e91627 (2023).

61. Suzuki, M. & Larkum, M. E. General Anesthesia Decouples Cortical Pyramidal Neurons. Cell 180, 666–676.e13 (2020).

62. Vangeneugden, J. et al. Activity in Lateral Visual Areas Contributes to Surround Suppression in Awake Mouse V1. Current Biology 29, 4268–4275.e7 (2019).

63. Turrigiano, G. G. The Self-Tuning Neuron: Synaptic Scaling of Excitatory Synapses. Cell 135, 422–435 (2008).

64. Turrigiano, G. G., Leslie, K. R., Desai, N. S., Rutherford, L. C. & Nelson, S. B. Activitydependent scaling of quantal amplitude in neocortical neurons. Nature 391, 892–896 (1998).

65. Ringach, D. L. & Malone, B. J. The operating point of the cortex: neurons as large deviation detectors. J Neurosci 27, 7673–7683 (2007).

66. Adesnik, H., Bruns, W., Taniguchi, H., Huang, Z. J. & Scanziani, M. A neural circuit for spatial summation in visual cortex. Nature 490, 226–231 (2012).

67. Kondo, S., Kikuta, K. & Ohki, K. Synaptic input architecture of visual cortical neurons revealed by large-scale synapse imaging without backpropagating action potentials elife (2025).

68. Yuste, R. & Denk, W. Dendritic spines as basic functional units of neuronal integration. Nature 375, 682–684 (1995).

69. Chen, T.-W. et al. Ultrasensitive fluorescent proteins for imaging neuronal activity. Nature 499, 295–300 (2013).

70. Müller, W. & Connor, J. A. Dendritic spines as individual neuronal compartments for synaptic Ca2+ responses. Nature 354, 73–76 (1991).

71. Schiller, J., Major, G., Koester, H. J. & Schiller, Y. NMDA spikes in basal dendrites of cortical pyramidal neurons. Nature 404, 285–289 (2000).

72. Chen, Y. et al. Thalamic activation of the visual cortex at the single-synapse level. Science 391, 1349–1354 (2026).

73. Weiler, S. et al. Orientation and direction tuning align with dendritic morphology and spatial connectivity in mouse visual cortex. Current Biology S0960982222002810 (2022).

74. Wilson, D. E., Whitney, D. E., Scholl, B. & Fitzpatrick, D. Orientation selectivity and the functional clustering of synaptic inputs in primary visual cortex. Nat Neurosci 19, 1003–1009 (2016).

75. Chen, X., Leischner, U., Rochefort, N. L., Nelken, I. & Konnerth, A. Functional mapping of single spines in cortical neurons in vivo. Nature 475, 501–505 (2011).

76. Scholl, B., Wilson, D. E. & Fitzpatrick, D. Local Order within Global Disorder: Synaptic Architecture of Visual Space. Neuron 96, 1127–1138.e4 (2017).

77. Kirkcaldie, M. T. K. Neocortex. in The Mouse Nervous System 52–111 (Elsevier, 2012). doi:10.1016/B978-0-12-369497-3.10004-4.

78. Guizar-Sicairos, M., Thurman, S. T. & Fienup, J. R. Efficient subpixel image registration algorithms. Opt. Lett. 33, 156 (2008).

79. Rossi, L. F., Harris, K. D. & Carandini, M. Spatial connectivity matches direction selectivity in visual cortex. Nature 588, 648–652 (2020).

80. Pachitariu, M. et al. Suite2p: Beyond 10,000 Neurons with Standard Two-Pho-ton Microscopy. bioRxiv (2016).

81. Cuntz, H., Forstner, F., Borst, A. & Häusser, M. The TREES toolbox--probing the basis of axonal and dendritic branching. Neuroinformatics 9, 91–96 (2011).

82. Brainard, D. H. The Psychophysics Toolbox. Spatial Vis 10, 433–436 (1997).

83. Stringer, C. et al. Extracting large-scale neural activity with Suite2p. bioRxiv (2026).

84. Dipoppa, M. et al. Vision and Locomotion Shape the Interactions between Neuron Types in Mouse Visual Cortex. Neuron 98, 602–615.e8 (2018).

85. Schröder, S. et al. Arousal Modulates Retinal Output. Neuron 107, 487–495.e9 (2020).

